# Evaluation of High-Affinity Monoclonal Antibodies and Antibody-Drug Conjugates by Homogenous Time-Resolved FRET

**DOI:** 10.1101/2024.08.05.606727

**Authors:** Harmon Greenway, Jin Wang

## Abstract

The rapid growth of therapeutic monoclonal antibodies demands greater accessibility to scalable methods of evaluating antigen binding. Homogenous TR-FRET is ideal for preliminary screening but has not been reported to assay these interactions due to their high-affinity and complex solution-phase kinetics. Here we report the development of a competition assay to rank-order the relative affinities of these drugs for a common antigen. The assay is compatible with automation, requires no modification of the analytes, and measures affinities as low as single-digit picomolar. We further demonstrate applications to inform the development of antibody-drug conjugates. The assay may aid discovery and manufacturing of therapeutic antibodies as a low-cost, high-throughput alternative to existing technologies.

---

**SCHEME 1:**
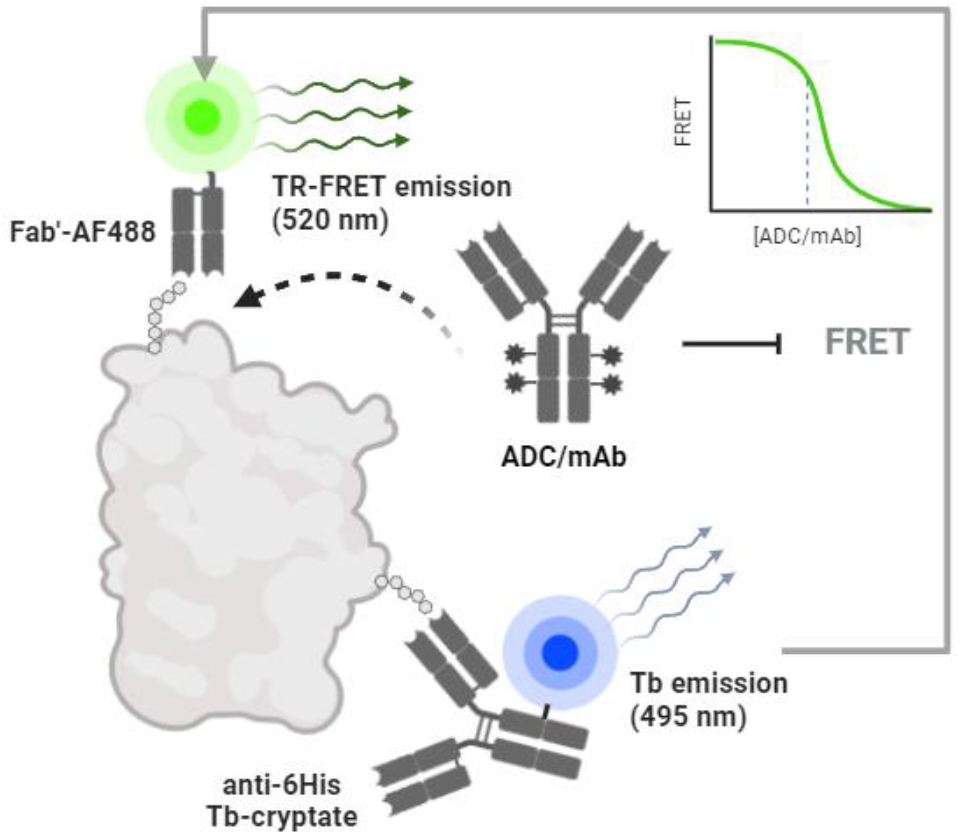
Competition TR-FRET assay format with Alexafluor-488 labeled IgG Fab displaced from Tb-labeled target protein by mAb-based therapeutic.^1^

Monoclonal antibodies (mAbs) and antibody-drug conjugates (ADCs) are therapeutic modalities with broad scope in the treatment of human diseases including cancer, autoimmune disorder, and infectious illness.^2,3,4^ These targeted drugs rely on specific, high-affinity interactions with antigens. Understanding target engagement is essential to drug development, and efficient methods to evaluate binding affinity are required for screening and quality control. The immune system has evolved an accelerated mechanism of affinity maturation and positive selection to identify high-affinity antibodies.^5^ This has been essential to the progress of mAb-based therapeutics, as in vitro binding assays present uncommon pitfalls and complex artifacts. The solution-phase kinetics of mAbs and antigens often defy analysis when ternary complexes are formed. Similarly, avidity effects preclude the flow of mAbs over a solid-phase.^6^ These interactions can be mitigated by immobilization of the antibody onto the surface of a biosensor chip or bead, but this requires additional chemistry and instrumentation. We describe an expedient method that leverages the competitive displacement of a fragment of antigen binding (Fab) from the recombinant extracellular domain of the target antigen in solution(Scheme 1). The assay offers an accessible “mix-and-read” approach to evaluate the binding affinity of mAbs and ADCs.

Homogenous TR-FRET is a fluorescence-based technique that is commonly used for high-throughput drug screening. It observes binding events by detecting the nanometer-scale proximity of two fluorophores.^7^ Förster resonance energy transfer is a non-radiative transfer of energy from an excited-state fluorescent donor to an acceptor fluorophore with suitable spectral overlap. The long fluorescence lifetime of the donor, typically a lanthanide chelate, enables time-resolved detection to eliminate background and scattered light.^8^ The high sensitivity and low signal-to-noise ratios of TR-FRET facilitate low sample requirements. The method is frequently used to study proteinprotein interactions but has not been reported to assay the antigen binding affinity of mAb-based therapeutics.

Antibodies produced by affinity maturation often demonstrate ‘tight-binding’ to their target antigens. Tight-binding is characterized by depletion of free ligand and occurs whenever the receptor concentration is greater than the equilibrium dissociation constant (K_D_). Michaelis-Menten kinetics cannot evaluate conditions of ligand depletion, as the model assumes the concentration of free ligand is fixed. The resulting error can be significant, particularly in low volume assays.^9^ Instead, tight-binding demands exact analytical equations. The K_D_ of binary receptor-ligand complex formation is correctly evaluated by the quadratic model reported by Morrison.^10^ (Figure 1A) The equilibrium inhibition constant (K_I_) values of two competing ligands can also be obtained with the cubic model reported by Zhi-Xin Wang.^11^ (Figure 1B) However, these models are merely reductive approximations of tight-binding in the presence of ternary complexes.

**FIGURE 1:**
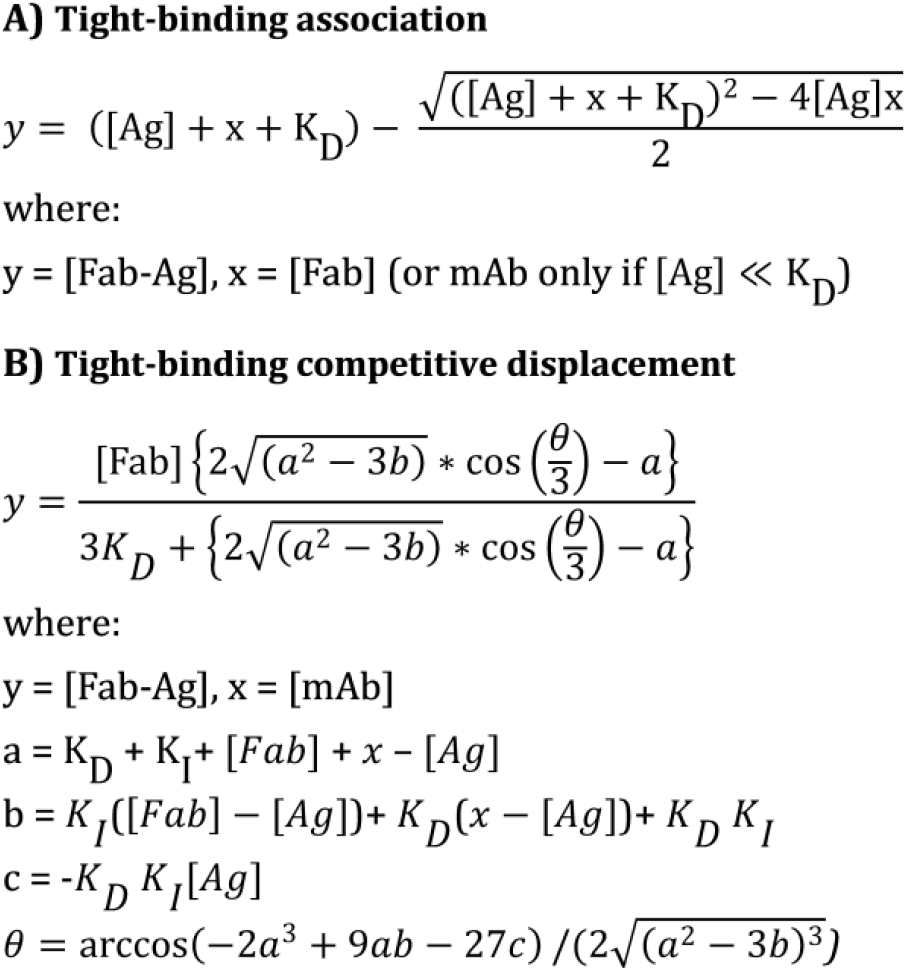
Equations for evaluating tight-binding kinetic parameters. (A) Morrison equation for association (B) Wang equation for competitive displacement titration. The equations do not account for multivalence and evaluate mAb-antigen binding by reductive approximation.

Immunoassays using mAbs are susceptible to an artifact known as the ‘high-dose hook effect’ or ‘antigen excess effect’ which manifests as a paradoxical decrease in analyte signal with increasing concentration.^12^ The artifact may be visually apparent as an inflection in association kinetics or the dose-response curve. This falsely-low response is driven by the formation of multivalent antibody-antigen complexes.^13^ When the hook effect is encountered, it is typically corrected by serial dilution.^14^ Unfortunately, the coincidence of tight-binding and multivalence is analytically intractable. As a classical three-body problem, no exact equation may be derived to account for the formation of ternary mAb-Ag complexes in ligand depletion regimes.^15^ Despite this limitation, we have generated significant insights into the behavior of these systems with computational models.

Assays of mAb-antigen binding may demand costly equipment or suffer from low throughput and high sample requirements. The affinities of mAb-based drugs are generally evaluated with surface-plasmon resonance (SPR), biolayer interferometry (BLI), or the kinetic exclusion assay (KinExA).^16, 17^ These techniques demand immobilization of the mAb to avoid avidity effects or measure free ligand. Apparent rate constants among these assays can vary by orders of magnitude due to slow dissociation rates, surface-based artifacts, or immobilization chemistries.^18^ The values reported by KinExA may better represent solution-phase rate constants, but this method has the lowest throughput.^19^ Observing the intrinsic kinetics of highaffinity mAbs remains a technical challenge. However, the correct rank-order and relative affinity is sufficient to inform screening and quality control applications.

This study highlights that high-affinity mAb-based therapeutics can be effectively rank-ordered by competition TR-FRET without immobilization. Our reported TR-FRET assay consistently discriminated mAbs of predictable relative affinities with single digit picomolar sensitivity. Advantages of this workflow include no prerequisite for labeling or modification of the mAb, low sample requirements, and full compatibility with automation. Alongside other methods, binding assays are useful to validate the specific activity of mAb-based therapeutics and may identify faults in quality control. We designed a simple set of experiments to investigate how inappropriate storage and handling of mAb proteins can negatively alter their binding affinity. Analysis of those experiments by competition TR-FRET confirmed that the appropriate conditions of manufacturing and storage are critical to ensure activity.

## RESULTS AND DISCUSSION

### Mass-balance models of bivalent tight-binding

An assessment of homogenous binding assays for high-affinity mAbs was conducted in MatLab Simbiology by deriving systems of ordinary differential equations with mass balance principles (Figures S10, S16).^20^ Although these systems have no analytical solutions, simulated doseresponse curves were produced with numeric solvers and evaluated by reductive approximation with the equations of Morrison or Wang (Figure 2). The plots compare the evaluated rate constants with the corresponding input values used to generate each simulation. In general, if the antigen concentration falls well below the K_D_ of the mAb-Ag interaction, then formation of the ternary complex is negligible and the apparent rate constants reflect the true values. However, when the antigen concentration exceeds the K_D_, formation of the ternary complex becomes significant, and the apparent constants deviate. The magnitude of this error depends on the degree of antigen excess and format of the assay.

**FIGURE 2:**
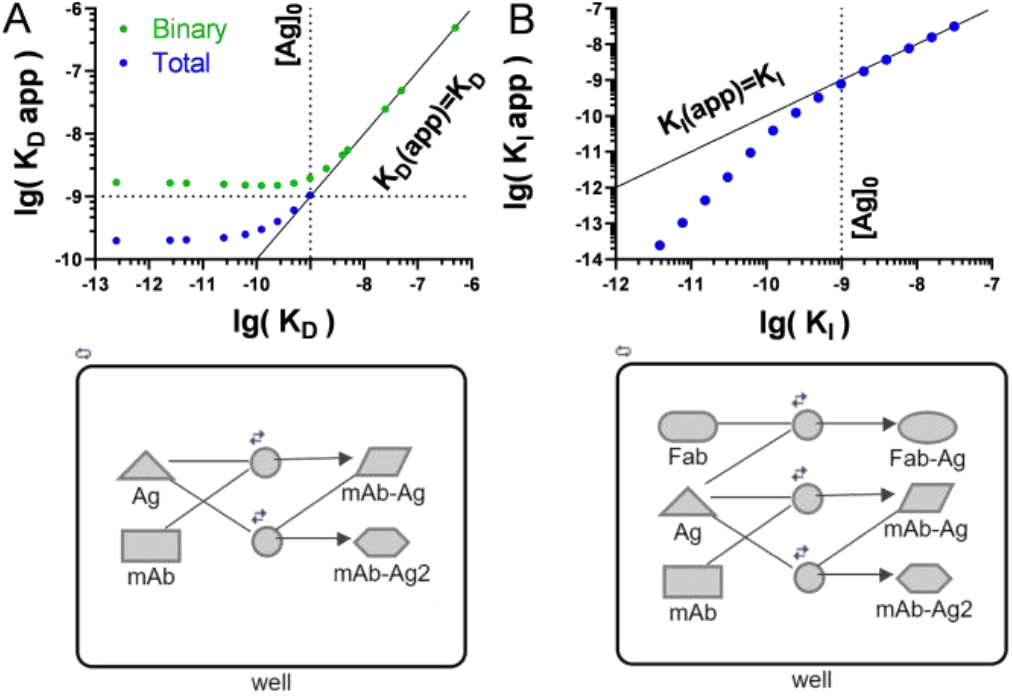
Comparison of rate constants from simulated direct and competition assays; apparent (y-axis) vs. input (x-axis) values. (A) Log-log plot of K_D_ values (Morisson) from mAb-Ag association simulations (top). Model of association (bottom). (B) Log-log plot of K_I_ values (Wang) from mAb displacement of Fab-Ag complex (top). Model of competition (bottom).

Direct assays in homogenous solution are particularly sensitive to the hook effect. The model of mAb-Ag association indicates that a direct binding assay will observe false equivalence among apparent affinities below the antigen concentration (Figure 2A). The trend occurs regardless of whether the K_D_ or EC_50_ is evaluated, and irrespective of whether the analyte is the totality of bound mAb or the binary complex alone. As the solution-phase K_D_ is often not known *a priori*, this implies a potential for type-I (false positive) or type-II (false negative) error in immunoassays. The limit of antigen excess is defined by the K_D_ of the mAb and antigen in the given conditions. Immunoassay development may benefit from highlighting this limit to prevent type I/II error. The model is sufficient to account for the widespread adoption of immobilization techniques in direct binding assays.

Competition assays are routinely used when an analyte signal cannot be observed. In contrast to direct assays, they evaluate affinity by measuring the displacement of another ligand by the analyte. The model of the competition format suggests that if a monovalent ligand is displaced from a receptor by disparate bivalent analytes, the false equivalence correlated with tight-binding in homogenous assays is not observed (Figure 2B). In this case, the rank-order of analytes is maintained and the potential for the type I/type-II errors associated with the hook effect is eliminated. The limit of antigen excess is not strictly defined by the K_D_ of the mAb in a competition assay and is also relative to the concentration and affinity of the competitor. Therefore, we performed a series of experiments to determine if competition TR-FRET reliably detects differences in the binding strength of high-affinity mAbs.

### Experimental design and evaluation of trastuzumab at 4ºC

The competition TR-FRET assay is performed in a 384-well plate and may use any suitable FRET pair to label a Fab or nanobody and corresponding antigen. The variable region may have an identical sequence to the analyte mAb or bind preferentially to a desirable epitope. In the following examples, a commercially available anti-6His terbium(Tb)-cryptate serves as a donor fluorophore to label the recombinant extracellular domain (ECD) of human HER2 antigen. The acceptor fluorophore is synthesized by conjugating Alexafluor-488 (AF488) maleimide to a Fab’ fragment. Each sample is purified prior to the assay, and concentrations are determined by UV spectroscopy. All pipetting is performed by an automated microfluidic dispenser. Analyte mAbs are diluted in series and used as titrants to a fixed concentration of the labeled Fab and antigen in triplicate test wells. The Fab’-AF488 tracer binds the Tb-labeled antigen and produces a FRET emission upon excitation of the donor fluorophore. The ratio of FRET emissions is proportional to the degree of specific binding between the Fab and antigen. Competitive displacement of the Fab from the antigen by the analyte inhibits the FRET signal and gives a dose-response. These interactions are both evaluated on the same plate to control for equilibration at measurement. Any non-specific background is recorded in control wells and subtracted from the signals. The rate constants are then determined by fitting the normalized dose-response curves by reductive approximation to exact analytical equations for binary complexes.

The affinity of a synthesized Fab’-AF488 and its parent trastuzumab for the HER2-ECD were evaluated by TR-FRET at 4ºC. The dose-response curves were fitted with the Morrison and Wang equations to determine the apparent K_D_ and K_I_ values, respectively. The association plots of Fab’-AF488 at concentrations 0-12.8 nM with HER2-ECD illustrate typical titration curves and each trace was fit by the Morrison equation with R^2^>0.99 (Figure 3A). The inhibition plots of trastuzumab at concentrations 0-128 nM with the Fab-HER2 complex were evaluated against the apparent K_D_ of the Fab and each trace was fit by the Wang equation with an R^2^>0.99 (Figure 3B). Equilibration was essentially complete after 18 hours at 4ºC. Although the apparent rate constants may be evaluated from a single measurement at equilibrium, the full equilibration curve was often monitored. A poor fit of this curve to a one-phase decay model may indicate variance related to biphasic kinetics or assay stability. The apparent dissociation constants of Fab’-AF488 from 2-40 h with 0.5 nM HER2-ECD antigen at 4ºC plateau at K_D_=10.4 pM (8.7-12 pM) and fit a one-phase decay model with R^2^ of 0.99. The apparent inhibition constants observed during equilibration of trastuzumab with 0.5 nM HER2-ECD antigen and 2 nM Fab’-AF488 at 4ºC plateau at K_I_=7.1 pM (4.7 pM-9.3 pM) and fit a one-phase decay model with R^2^ of 0.97. The decay of rate constants over time reflects the expected behavior of stable equilibration and 1:1 binding interaction.

**FIGURE 3:**
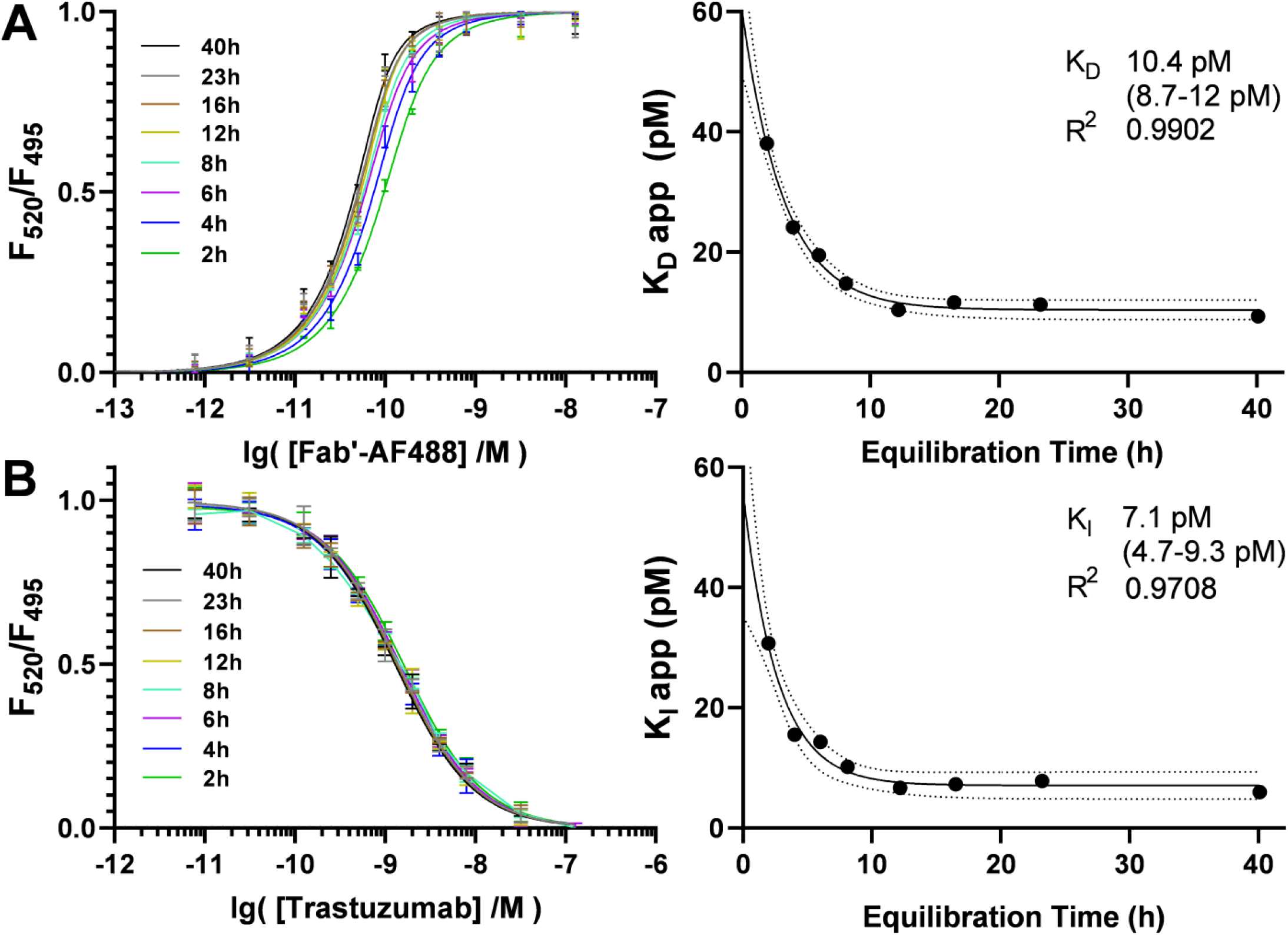
(A) Evaluation of Fab’-AF488 K_D_. Association curves observed by normalized FRET ratio on titration of Tb-labeled HER2-ECD with Fab’-AF488 from 2-40 hours at 4ºC (left) and equilibration of the apparent K_D_ values evaluated with the Morrison equation and fit with one-phase decay. (right) (B) Dissociation curves observed by normalized FRET ratio of Fab’-AF488 and Tb-labeled HER2-ECD from 2-40 after titration with trastuzumab (left) and equilibration of apparent K_I_ values evaluated with the Wang equation and fit with one phase decay. (right)

### Investigation of reagent concentration regimes

A TR-FRET binding assay should utilize the lowest antigen concentration that provides good signal-to-noise for the attached fluorophore on the instrument. A fundamental difference between K_I_ and IC_50_ values is that a rate constant should not vary with receptor concentration. However, antigen excess in a binding assay may alter the apparent rate constants of mAb analytes due to the hook effect. Therefore, the effect of antigen concentration was studied to determine the limit at 4ºC. A series of competition assays were conducted between trastuzumab and 1 nM Fab’-AF488 at concentrations of HER2-ECD from 250 pM-4 nM (Figure 4A). After an 18-hour incubation period, the doseresponse curves were evaluated. Assays conducted at 250 pM, 500 pM, and 1 nM HER2 all observed a K_I_ of trastuzumab at 10.8 pM ± 0.03 pM at 4ºC, validating this regime of antigen concentration for use in the assay. The apparent K_I_ is observed to decrease slightly with antigen concentrations above 1 nM, consistent with antigen excess.

**FIGURE 4:**
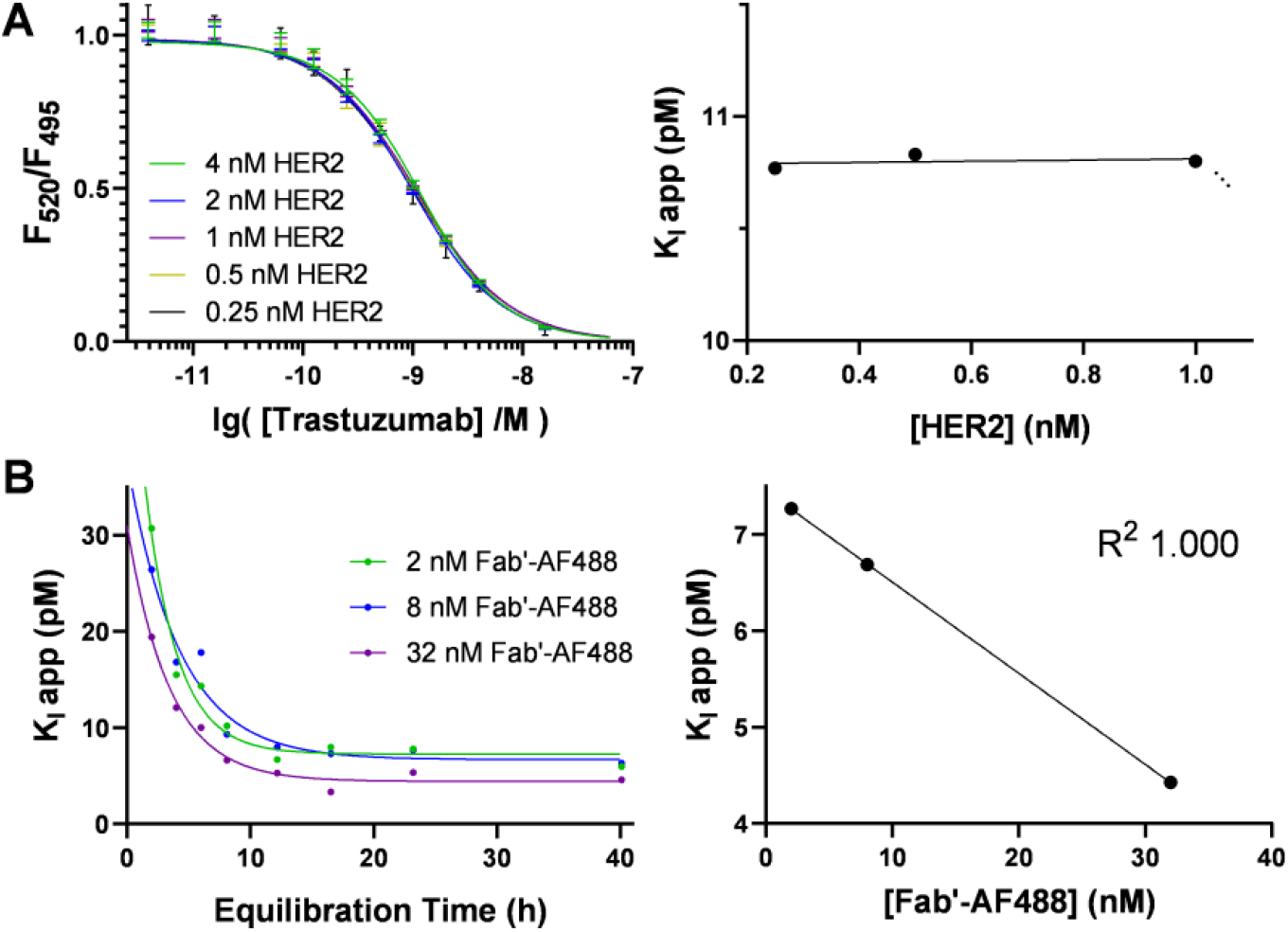
(A) Effect of antigen concentration. Dissociation curves of competition assays with increasing fixed antigen concentration after 18h incubation at 4ºC (left) and resulting K_I_ values corresponding to 0.25-1 nM HER2. (right) (B) Effect of tracer concentration. Equilibration curves of competition assays with increasing fixed concentrations of Fab’-AF488 tracer (left) and fit of resulting K_I_ values (right).

The tracer concentration of the TR-FRET assay should be minimally sufficient to saturate the antigen. An excess of labeled competitor may inhibit the formation of ternary complexes but also increases the concentration of fluorophore. Models indicate that increasing the concentration of the competitor will rescue a ‘hooked’ assay and otherwise have no effect on the result (Figure S21). However, photophysical interferences may attenuate the response signal, such as donor quenching or inner filter effects.^21^ The effect of increasing concentrations of the Fab’-AF488 tracer on the apparent K_I_ of trastuzumab was further studied. Competition assays were conducted between trastuzumab and 0.5 nM HER2-ECD at tracer concentrations of 2 nM, 8 nM, and 32 nM. The apparent K_I_ in each assay was evaluated periodically from 2-40 hours and fit to a one-phase decay model to determine the values at equilibrium (Figure 4B). The apparent K_I_ of trastuzumab at 4ºC decreases with increasing tracer concentration over the range 7.3-4.4 pM. Linear regression of K_I_ values with the corresponding Fabtracer concentrations gives a perfect R-squared value of 1.000. The K_I_ observed at a tracer concentration of 2 nM falls 0.2 pM under the y-intercept of the slope. These results reflect the precision of the instrument and may estimate the significance of photophysical interferences in evaluation of apparent K_I_ values.

### Applications of TR-FRET in quality control of ADCs

Among our other published applications of TR-FRET in drug discovery^22^, the assay was developed as a means of quality control for antibody-drug conjugates. The potency of an ADC relies on its ability to engage the target, and this may be compromised at any stage of production or storage. Affinity measurements may supplement good manufacturing practices by identifying inconsistent binding of samples at each stage. To evaluate the suitability of TR-FRET for this application, we subjected trastuzumab to various conditions expected to compromise the quality of antibodies during the synthesis of ADCs. These also provided a means to create a predictable rank-order of binding affinity among a set of antibody samples. We studied the effect of short-term exposure to organic solvents, as well as freezing temperature and repeated freeze/thaw cycles. We investigated the consequences of breaking disulfide bonds by reduction, a necessary step in the production of cysteine-linked ADCs. The stoichiometry of conjugation for trastuzumab emtansine, a lysine-linked ADC, was also explored. The competition TR-FRET assay illustrated proportional trends among the affinities of treated mAbs which correlate these conditions with a potential to reduce binding affinity when optimized parameters are exceeded. The results underscore the importance of adhering to best practices in ADC synthesis.

### Disulfide reduction and cysteine-linked ADCs

The conjugation of cysteine-linked ADCs requires reduction of interchain disulfide bonds in the hinge region of the antibody. The resulting thiol may be treated with maleimides or other Michael acceptors to attach a drug payload.^23^ An excess of the reducing agent may be used to increase drug loading. However, the reagents employed are not generally regiospecific to the hinge, and over-reduction may disrupt structurally relevant disulfides. For this reason, some protocols cite subsequent attempts to re-bridge disulfide bonds with dehydroascorbic acid.^24^ The preferred reducing agent is tris(2-carboxyethyl)phosphine hydrochloride (TCEP), which acts with near stoichiometric efficiency. An IgG1 antibody contains 16 disulfide bonds, of which 4 are in the hinge region.^25^ To investigate the effect of over-reduction on the apparent K_I_ of mAbs, we exposed samples of trastuzumab to increasing concentrations (0-32 eq.) of TCEP under similar conditions employed in ADC synthesis (Figure 5A). No significant loss of binding activity was observed after treatment with up to 8 equivalents TCEP relative to the control. Treatments of 8-32 equivalents resulted in an apparent exponential increase of K_I_ from 7-23 pM which may indicate denaturation. The assay illustrates a clear trend among the apparent K_I_ values and demonstrates that a large excess of TCEP compromises binding affinity.

**FIGURE 5:**
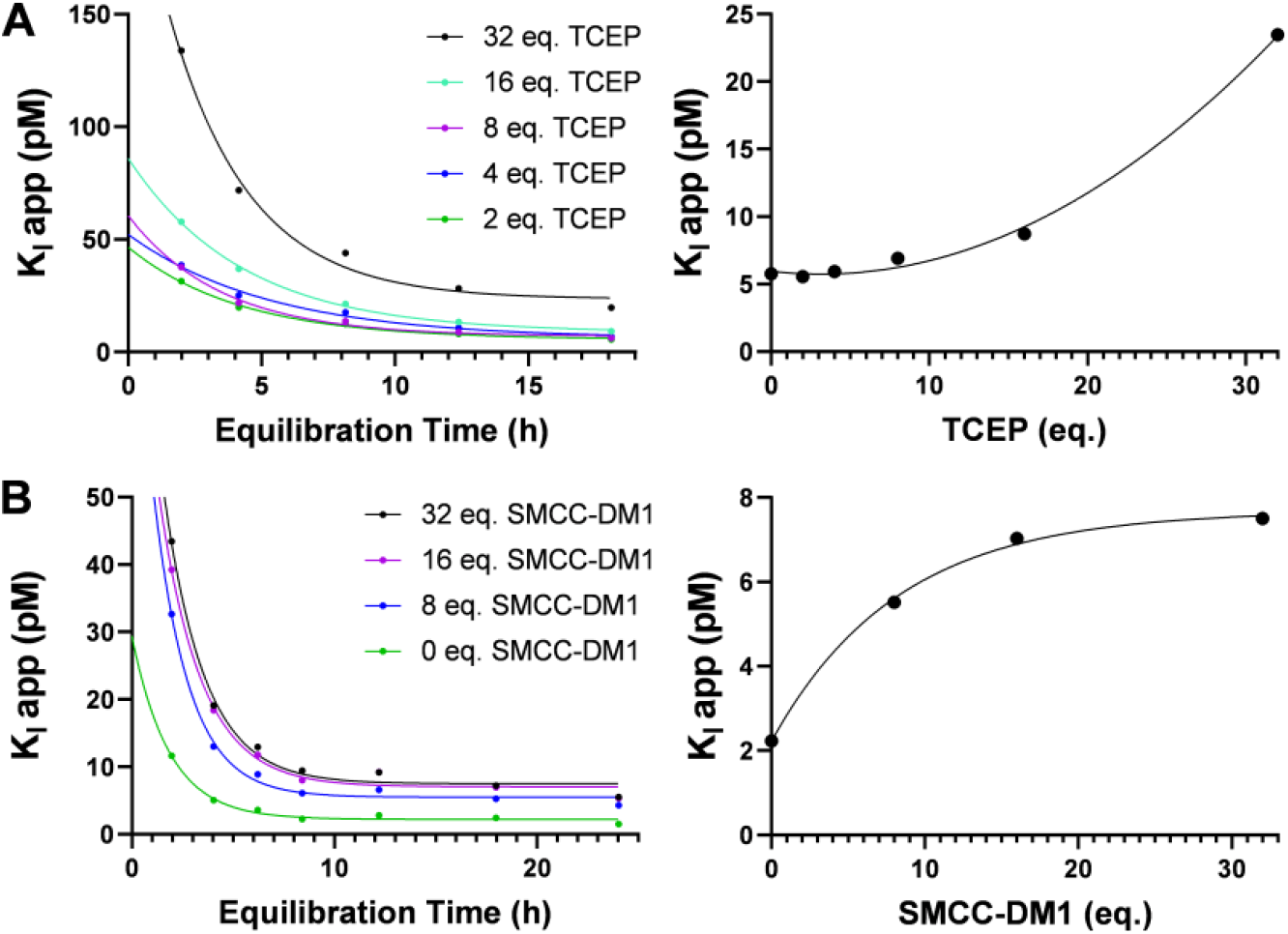
(A) Equilibration curves of apparent K_I_ values from 2-18h for trastuzumab samples reduced with increasing equivalents of TCEP (right) and plot of equilibrium K_I_ with reduction stoichiometry (left). (B) Equilibration curves of apparent K_I_ values from 2-24h for trastuzumab samples conjugated with increasing equivalents of smcc-DM1 (right) and plot of equilibrium K_I_ with conjugation stoichiometry (left).

### Conjugation stoichiometry and lysine-linked ADCs

The conjugation of lysine-linked ADCs is conducted by acylation of protonated lysine residues on the solvent-exposed surface of the antibody. Attachment of multiple drug molecules results in a heterogenous distribution which significantly influences the pharmacokinetics and activity of the conjugate.^26^ A stoichiometric excess is often employed to increase the drug-to-antibody ratio (DAR). However, the average DAR may be limited by the hydrophobicity of the drug payload, as high-DAR populations can form aggregates which are removed during purification. The remaining distribution may not represent an intact population of the same DAR. To study the effect of conjugation stoichiometry on apparent binding affinity, a series of trastuzumab emtansine (T-DM1) conjugates were synthesized at 40ºC with moderate to excessive amounts of smcc-DM1. **Caution!** Cytotoxic drugs can be hazardous in small quantities. The average DAR of conjugates from 8, 16 and 32 equivalents was estimated by UV spectroscopy as 2.1, 3.2, and 2.7, respectively. Treatment with smcc-DM1 resulted in an apparent logarithmic increase from 2-8 pM in the K_I_ of a trastuzumab sample that may illustrate a saturation curve (Figure 5B). The assay does not relate DAR with the K_I_ of emtansine conjugates or evaluate solvent-exposed emtansine distributions. However, the correlation implies that using excessive drug-linker to achieve a target DAR may reduce the affinity of some lysine-linked ADCs, particularly when the payload is hydrophobic.

### Denaturing conditions in the conjugation buffer

Although many buffer conditions are employed in the conjugation of ADCs, an organic component is often required to dissolve hydrophobic drug-linkers. This is typically a polar, aprotic solvent such as dimethylformamide (DMF) or dimethylsulfoxide (DMSO). The tertiary structure of a protein is optimized such that hydrophilic residues are solvent-exposed and hydrophobic residues are buried in the interior. Exposure to organic solvents may disrupt this arrangement. To compare the effects of these solvents on the apparent affinity of mAbs, we exposed samples of trastuzumab to buffers containing increasing proportions of DMF and DMSO for a two-hour period (Figure 6A). The apparent binding affinities of all samples containing up to 10% organic solvent were equivalent to the control. Surprisingly, the K_I_ of samples exposed to 25-50% DMSO also remained constant. In contrast, exposure to 25-50% DMF resulted in an apparent exponential increase of K_I_ values from 10-36 pM which may indicate denaturation. These results do not imply that trastuzumab does not rapidly aggregate in high concentrations of DMSO. We infer that the tertiary structure of the monomeric fraction recovered or remained intact. The result is consistent with the relative polarity of the solvents, and this may suggest that DMSO is superior for use in bioconjugation. The results highlight that conjugation buffers should contain the minimum volume of organic solvent required to dissolve the reagents.

**FIGURE 6:**
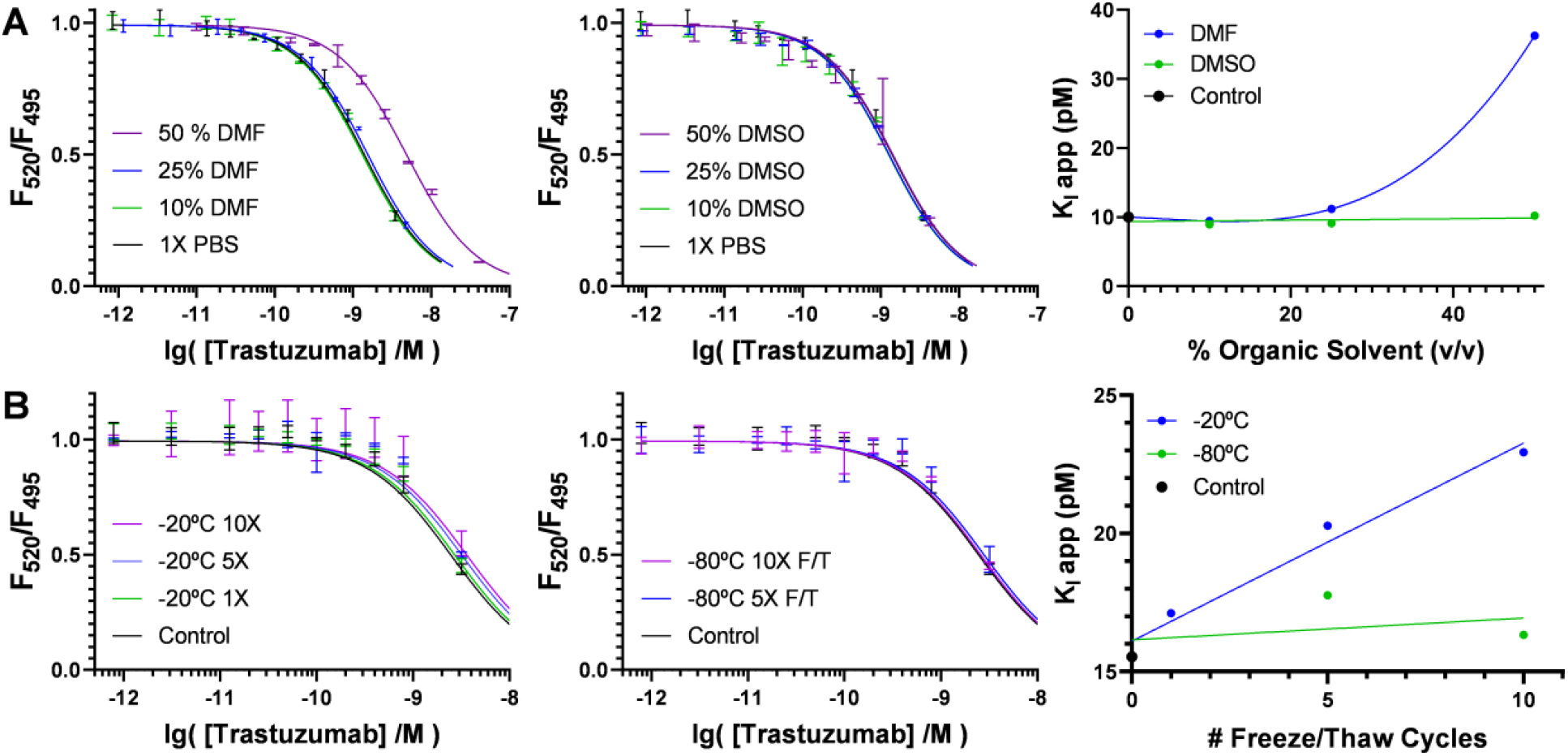
(A) Dissociation curves observed by normalized FRET ratio of Fab’-AF488 and Tb-labeled HER2-ECD 24h after titration with trastuzumab analytes exposed to buffers containing DMF (left) or DMSO (middle) and plot of apparent K_I_ with the proportion in each buffer (right). (B) Dissociation curves observed by normalized FRET ratio of Fab’-AF488 and Tb-labeled HER2-ECD 24h after titration with trastuzumab analytes exposed to 0-10X freeze/thaw cycles at −20ºC (left) or −80 ºC (middle) and plot of apparent K_I_ with freezing temperature (right).

### Denaturing conditions in cold storage

Antibodies, like other proteins, are sensitive to freeze-thaw cycles. The shelf-life of IgG antibodies at 4ºC may vary from several weeks to many years.^27^ It has been reported that trastuzumab is stable at 2-8ºC for at least 28 days.^28^ However, ensuring the efficacy of therapeutic mAbs and ADCs requires freezing. The velocity at which they are frozen is significant, as the tensile stress of ice crystal formation induces changes in their native conformation. Although the best practice is freezing with liquid nitrogen followed by lyophilization, antibodies are often stored in freezers at - 20ºC or −80ºC in laboratory settings. To evaluate the effect of freezing velocity, we subjected aliquots of 5 mg/mL trastuzumab in PBS to up to 10 freeze-thaw cycles at these temperatures (Figure 6B). The K_I_ values observed for samples frozen at −20ºC exhibited an apparent linear increase with an R^2^ value of 0.974. The relative slopes of regression reflect that the binding affinities of mAbs recovered from cold storage are inversely proportional to the temperature of freezing. The experiment emphasizes that antibodies must not be frozen at −20ºC without cryoprotectants and should be aliquoted prior to freezing at any temperature to avoid excessive freeze/thaw cycles.

### Conclusions

These experiments collectively establish that competition-format TR-FRET effectively evaluates the rank-order and relative binding strength of high-affinity mAbs. This topic may be poorly understood, as the coincidence of tight-binding and multivalence is analytically intractable. Homogenous immunoassays using mAbs are often susceptible type-I/type-II errors due to the hook effect. Our models demonstrate that this is a function of antigen concentration and binding affinity, where the equilibrium dissociation constant can represent an inflection point between valid and invalid results in direct assays. However, these models also indicate that the competition-format assay is not susceptible to these errors, as the rank-order of analytes was consistently maintained when a high-affinity competitor is present to mitigate the formation of multivalent mAb-Ag complexes.

An antigen concentration regime should still be defined against the highest affinity mAb at the outset of the experiment to avoid error in the apparent rate constants. Our assay of trastuzumab demonstrated femtomolar precision across a range of antigen concentrations exceeding the apparent K_D_/K_I_ values at 4ºC. However, the method permits evaluation of K_I_ values lower than the antigen concentration only when a suitable competitor is used. Highaffinity antibody fragments or nanobodies are well-suited to this requirement. Photophysical interference is a common source of error in ratiometric fluorimetry which may be estimated by defining a concentration regime for the acceptor fluorophore. Assisted by an automated microfluidic dispenser, we demonstrated the precision of the instrument in the equilibrium inhibition constants of assays with increasing concentrations of the labeled-Fab. The results of HER2 and Fab’-AF488 concentration studies each validate the selected conditions and exclude significant error from these sources.

Rate constants may vary with temperature, and this assay required low temperature to remain stable over an equilibration period of 18 hours. It was demonstrated to remain stable for up to 40 hours at 4ºC. The slow dissociation rates and long equilibration times that characterize high-affinity mAbs are known challenges to the estimation of kinetic parameters by SPR.^29^ Although precise equilibration curves were illustrated by the apparent rate constants obtained by periodic measurement, only the endpoint data is required and may be collected after incubation. The affinity of trastuzumab for the HER2-ECD at 20ºC was previously reported by SPR with values ranging from 100 pM-9.4 nM, and these reports often agree on a K_D_ ∼0.5 nM for the Fab fragment.^30, 31,32^ It is typical for solution-phase assays of mAbs to observe higher affinities than surface-based techniques.^17,18^ However, the rank-order is sufficient to inform most drug development applications. An investigation of conditions relevant to the manufacturing and storage of antibody-drug conjugates demonstrates that affinity is a significant diagnostic of quality for therapeutic anti-bodies. Trastuzumab samples were treated by a series of harsh conditions which progressively deviated from optimized protocols. In each assay, a plot of the observed K_I_ values for the treated samples reflects a clear and proportional reduction of binding affinity. The examples are instructive and emphasize the need for optimized protocols to preserve the activity of the parent antibody.

The homogeneous, solution-phase format of the assay is more amenable to automation than existing techniques and is highly scalable. The advantage of the “mix-and-read” approach to screening is the lack of sample preparation requirements. Immobilization techniques demand additional chemistry and may limit throughput when each mAb analyte must be fixed to a surface. Label-free assays often have strict sample purification requirements and are not generally compatible with cell lysates. Analytical methods which are both adaptable to high-throughput screening and capable of evaluating low-picomolar affinities represent a largely unmet need in biologics drug development.^33^ The equipment and materials necessary to conduct TR-FRET are widely available to meet the rapidly growing demand for antibody research. The assay could be adapted for epitope binning^34^ studies or used to compare the results of in vitro affinity maturation techniques. Competitive TR-FRET may serve as a preliminary screen for the discovery of mAbs and can assist manufacturing of mAb-based therapeutics as an expedient diagnostic tool.

## SUPPORTING INFORMATION

Kinetic simulation results and systems of ordinary differential equations describing assay models; Synthesis and characterization of trastuzumab Fab’-AF488; General protocol for conducting the assay and curve fitting; Protocols for all experiments and binding curves with fitting for the corresponding assays

## AUTHOR INFORMATION

### Authors

Harmon Greenway - Verna and Marrs McLean Department of Biochemistry and Molecular Pharmacology, Baylor College of Medicine, Houston, Texas 77030, United States;

### Author Contributions

J.W. contributed to the conception and design of this research. H.G. contributed to the conception, design, and implementation of this research and wrote the manuscript. All authors contributed to manuscript revision, read, and approved the submitted version.

### Notes

J.W. is the co-founder of Chemical Biology Probes, LLC.

J.W. has stock ownership in CoRegen Inc and serves as a consultant for this company.

## ACKNOWLEDGEMENTS

The research was supported in part by National Institute of Health (R01-CA268518, and R01-CA250503 to J.W.), Cancer Prevention & Research Institute of Texas (CPRIT, RP220480 to J.W.), and the Michael E. DeBakey, M.D., Professorship in Pharmacology (to J.W.)., and Dr. Timothy Palzkill for use of FPLC equipment.

## ABBREVIATIONS

FRET: Forster resonance energy transfer
TR-FRET: timeresolved FRET
mAb: monoclonal antibody
Fab: fragment of antigen binding
F(ab’)2: two Fab fragments bound by disulfide from the hinge
Fab’: a Fab fragment that retains thiol from the hinge
ADC: antibody-drug conjugate
SPR: surface-plasmon resonance
BLI: biolayer interferometry
KinExA: kinetic exclusion assay
K_D_: equilibrium dissociation constant
K_I_: equilibrium inhibition constant
IC_50_: half-maximal inhibitory concentration
Ag: antigen
Tb: terbium
HER2: human epidermal growth factor receptor 2
ECD: extracellular domain
AF488: Alexafluor 488
TCEP: tris(2-carboxyethyl)phosphine hydrochloride
DAR: drug-to-antibody ratio
T-DM1: trastuzumab emtansine conjugate
smcc-DM1: emtansine
DMF: dimethylformamide
DMSO: dimethylsulfoxide
NaOAc: sodium acetate
DTPA: diethy-lenetriaminepentaacetic acid
EDTA: ethylenediamine tetraacetic acid
FPLC: fast protein liquid chromatography
Tris: Tris(hydroxymethyl)aminomethane
BSA: bovine serum albumin
PBS: phosphate-buffered saline
MWCO: molecular weight cut-off

## For Table of Contents Use Only

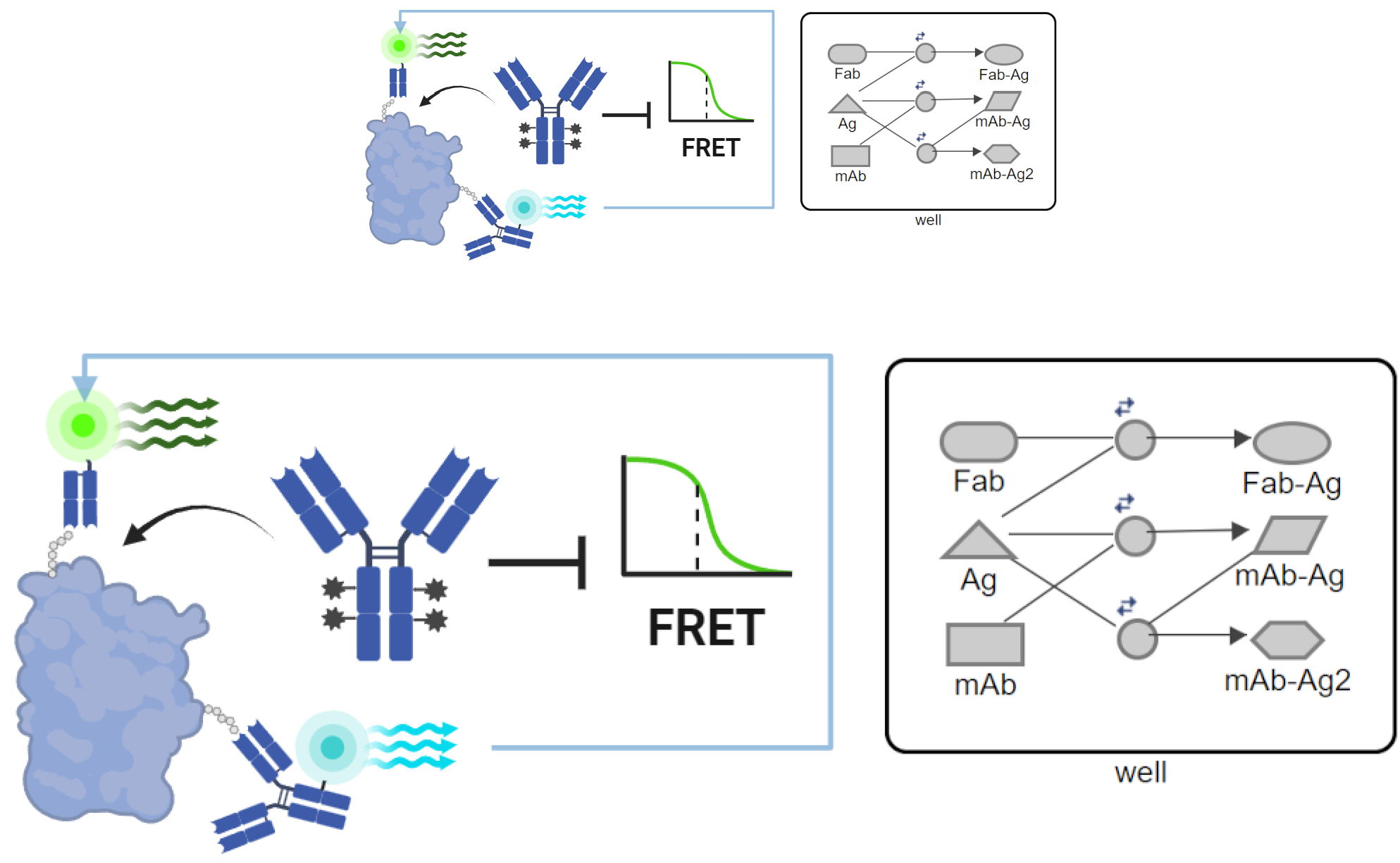

