## Supplemental Info for "Evaluation of High-Affinity Monoclonal Antibodies and Antibody-Drug Conjugates by Homogenous Time-Resolved FRET"

#### Supporting Information

Section 1. Determining Binding Affinity Constants from Concentration Response Data……………………………..S1

A. Analytical Evaluation of $K_D$ for Tight-Binding Ligands………………………………………………………S2

B. Mass-Balance Model of Tight-Binding Association………………………………………………………S2

C. Analytical Evaluation of $K_I$ for Competitive Tight-Binding Inhibitors…………………………………S3

D. Mass-Balance Model of Competitive Displacement for Tight-Binding Inhibitors……………………S4

Section 2. Bivalent Binding and Ternary Complex Equilibria in Conditions of Ligand Depletion…………………S5

A. Mass-Balance Model of Tight-Binding Ternary Complex Formation…………………………………S6

B. Analytical Intractability for Bivalent, Tight-Binding…………………………………………………S7

C. Mass-Balance Model of Bivalent, Tight-Binding Competitive Displacement…………………………S9

Section 3. Design of Competitive Displacement TR-FRET Assay for Evaluating Antibody Affinity……………..S11

A. Schematic Overview……………………………………………………………………………………S11

B. Synthesis of Fab'-Alexafluor488 TR-FRET Tracer…………………………………………………S12

C. Evaluating Drug-to-Antibody Ratio / Degree of Labeling by UV Spectroscopy………………………S13

D. General Protocol for Conducting Homogenous TR-FRET Assay………………………………………S14

E. Curve Fitting…………………………………………………………………………………………S15

Section 4. Evaluation of Concentration Regimes……………………………………………………………………S16

A. Effect of HER2-ECD Concentration in Apparent $K_I$ Evaluation by TR-FRET…………………………S16

B. Effect of Tracer Concentration on Apparent $K_I$ Evaluation by TR-FRET ……………………………S17

Section 5. Evaluation of Conditions Relevant to Production of Antibody-Drug Conjugates………………………S22

A. Effect of Solvent Exposure on the Binding Affinity of Trastuzumab……………………………………S22

B. Effect of Freeze/Thaw Cycles on the Binding Affinity of Trastuzumab…………………………………S24

C. Effect of TCEP Reduction on the Binding Affinity of Trastuzumab……………………………………S25

E. Synthesis and Effect of Stoichiometry on Apparent $K_I$ of Trastuzumab-DM1 ADCs…………………S31

Section 6. Supplementary References………………………………………………………………………………S35

#### Section 1. Determining Binding Affinity Constants from Dose-Response Data

To evaluate the binding affinity of monoclonal antibodies, it is useful to first review the equations with which binding constants are obtained in the general case. Under a normal binding regime, the dissociation constant ($K_D$) of a single receptor-ligand interaction may be obtained from a dose-response curve with the Michaelis-Menten equation (S1). The $K_D$ may be interpreted as the quotient of the rates of dissociation ($k_{off}$) and association ($k_{on}$) or as the ratio of free and bound species in solution. (S2) A competitive inhibitor of such a binding interaction may then be evaluated with the Cheng-Prusoff equation (S3), which relates the half-maximal inhibitory concentration to the inhibition constant ($K_I$) and concentration of labeled ligand.

$$B = (B_{max} * [L])/(K_D + [L]) \tag{S1}$$

$$K_D = k_{off}/k_{on} = ([R][L])/([RL]) \tag{S2}$$

$$K_I = IC_{50}/\ (1 + [S]/[K_M]) \tag{S3}$$

However, application of Michaelis-Menten kinetics requires the 'free ligand assumption', which implies that the activity of free ligand in solution remains approximately constant. It is often the case that monoclonal antibodies (mAbs) exhibit "tight-binding" to their target antigens, and this is characterized by depletion of free ligand. As the observed rate at any time point depends on the concentration of free ligand, these rates may vary if a significant portion of ligand becomes bound in the receptor-ligand complex. Ligand depletion is characteristic of tight-binding concentration regimes and will occur whenever the receptor concentration exceeds the $K_D$ of the ligand. Tight-binding and the condition of ligand depletion demand exact analytical solutions.

##### A. Analytical Evaluation of $K_D$ for Tight-Binding Ligands

An exact equation modeling the association of a receptor-ligand pair may be derived following the quadratic velocity equation published by Morrison. As the free concentration of any species in solution may be expressed as a function of its total and bound concentration (S4, S5), substitution of these expressions into the definition of $K_D$ (S6, S7) permits cross-multiplication (S8, S9) and expression as a quadratic formula which relates the $K_D$ to the dose-response curve. (S10) Unlike the Michaelis equation, the quadratic equation is always explicitly correct for binary receptor-ligand complexes.

$$[R] = [RL] + [R]f \quad (S4)$$

$$[L] = [RL] + [L]f \quad (S5)$$

$$K_D = ([R]f * [L]f)/[RL] \quad (S6)$$

$$K_D = ([R] - [RL])([L] - [RL])/([RL]) \quad (S7)$$

$$0 = ([R] - [RL])([L] - [RL]) - K_D[RL] \quad (S8)$$

$$0 = [RL]^2 - ([R] + [L] + K_D)[RL] + [R][L] \quad (S9)$$

$$[RL] = ( ([R] + [L] + K_D) - \sqrt{([R] + [L] + K_D)^2 - 4[R][L]} )/2 \quad (S10)$$

where; $[L]f$ = free ligand concentration

$[R]f$ = free receptor concentration

#### B. Mass-Balance Model of Tight-Binding Receptor-Ligand Association

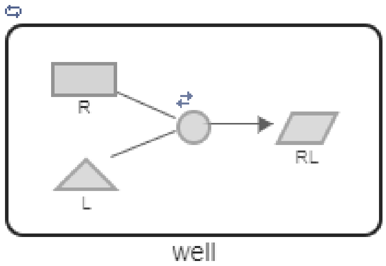


$$\frac{d[L]}{dt} = k_{off}[RL] - k_{on}[R][L]$$

$$\frac{d[R]}{dt} = k_{off}[RL] - k_{on}[R][L]$$

$$\frac{d[RL]}{dt} = k_{on}[R][L] - k_{off}[RL]$$

*Figure S1:* Graphical representation of receptor-ligand pair interaction in which a reversible association occurs.

*Figure S2:* System of ODEs representing a receptor-ligand pair association.

The validity of equation S10 to determine the dissociation constant of a receptor-ligand interaction can be demonstrated with simulated data. Following mass-balance principles, MatLab Simbiology software was used to generate an ODE-based model representing a titration of increasing concentrations of ligand to a fixed concentration of receptor. A graphical model of the interaction was produced (Figure S1) and a system of ODEs was calculated for the model. (Figure S2) The numeric solver (ODE15s) simulated a plot of the data with the relevant rate constants. (Figure S3) In this case, the values of $k_{on}$ = 7.1E5 and $k_{off}$ = 3.5E-4 ($K_D$=4.926E-10) were used to generate the simulation, and these values were used as a baseline in subsequent experiments unless otherwise specified. The equilibrium concentrations of the receptor-ligand complex corresponding to each concentration of ligand were then plotted in GraphPad Prism and fit to equation S10. As may be observed in this example, evaluation of the curve with the Morrison equation perfectly recapitulates the dissociation constant used to produce the simulation.

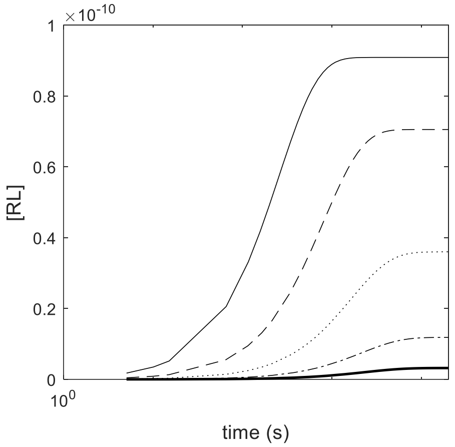


*Figure S3:* Semi-log plot of simulated of receptor-ligand association produced with $K_D$ = 4.9296E-10.

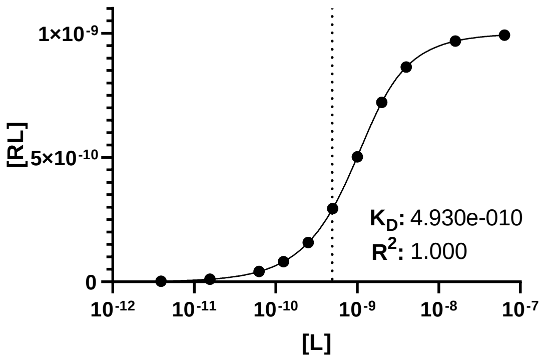


*Figure S4:* Semi-log plot of receptor-ligand association data from simulation with Equation S10.

#### C. Analytical Evaluation of $K_I$ for Tight-Binding Competitive Inhibitors

An exact analytical expression for competitive displacement of two tight-binding ligands (A and B) for a receptor (R) has been derived by Wang. In summary, as equations S11, S12, and S13 follow from mass-balance principles, and substitution of equation S12 and S13 into equation S11, followed by rearrangement, yields the cubic equation S14. Wang et. al determined that equation S14 has only one significant root which is equation S15. An exact expression for the dose response of either species is therefore given by the set of equations in S16. The rate constant of either species may be determined, if the other is known, from measurable parameters by curve fitting software without further derivation. The equation correctly accounts for ligand depletion in most competitive displacement titrations.

$$[R]_0 = [R] + [RA] + [RB] \quad (S11)$$

$$[RA] = ([R][A_0])/(K_A + [R]) \quad (S12)$$

$$[RB] = ([R][B_0])/(K_B + [R]) \quad (S13)$$

$$0 = [R]^3 + a[R]^2 + b[R] + c \quad (S14)$$

where:

$$a = K_A + K_B + [A_0] + [B_0] - [R]_0$$

$$b = K_B([A_0] - [R]_0) + K_A([B_0] - [R]_0) + K_A K_B$$

$$c = -K_A K_B [R]_0$$

$$[R] = -\frac{a}{3} + \frac{2}{3}\sqrt{(a^2 - 3b)} + \cos\left(\frac{\theta}{3}\right) \quad (S15)$$

where: $\theta = \arccos(-2a^3 + 9ab - 27c)/(2\sqrt{(a^2 - 3b)^3})$

$$[RA] = [A_0]\{2\sqrt{(a^2 - 3b)} * \cos\left(\frac{\theta}{3}\right) - a\}/(3K_A + \{2\sqrt{(a^2 - 3b)} * \cos\left(\frac{\theta}{3}\right) - a\}) \quad (S16)$$

$$[RB] = [B_0]\{2\sqrt{(a^2 - 3b)} * \cos\left(\frac{\theta}{3}\right) - a\}/(3K_B + \{2\sqrt{(a^2 - 3b)} * \cos\left(\frac{\theta}{3}\right) - a\})$$

##### D. Mass-Balance Model of Competitive Displacement for Tight-Binding Inhibitors

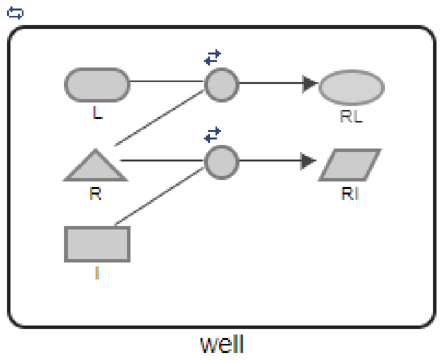


*Figure S5:* Graphical representation of a competitive displacement titration in which a receptor may be bound to a ligand or inhibitor.

$$\frac{d[L]}{dt} = k_{off}[RL] - k_{on}[R][L] \qquad \frac{d[I]}{dt} = k_r[RI] - k_f[R][I]$$

$$\frac{d[RL]}{dt} = k_{on}[R][L] - k_{off}[RL] \qquad \frac{d[RI]}{dt} = k_f[R][I] - k_r[RI]$$

$$\frac{d[R]}{dt} = k_{off}[RL] - k_{on}[R][L] + k_r[RI] - k_f[R][I]$$

*Figure S6:* System of ODEs representing a competitive displacement titration of two tight-binding inhibitors.

The validity of equation S16 to determine the inhibition constant of a competitive displacement titration can be demonstrated with simulated data. MatLab Simbiology software was used to generate ODE-based models following mass-balance principles, demonstrating the titration of increasing concentrations of inhibitor to a fixed concentration of receptor and ligand. A graphical model of the interaction was produced (Figure S5) and a system of ODEs was calculated for the model. (Figure S6) The binding constant $K_D$=$K_I$=4.9296E-10 was used to generate the data. A numeric solver (ODE15s) within MatLab simulated a plot of the data. (Figure S7) The equilibrium concentrations of the receptor-ligand complex corresponding to each concentration of inhibitor was then plotted in GraphPad Prism and fit to Supplementary Equation 10. As can be observed in this example, the curve-fitting with the Wang equation reproduces the inhibition constant used to generate the simulation. (Figure S8)

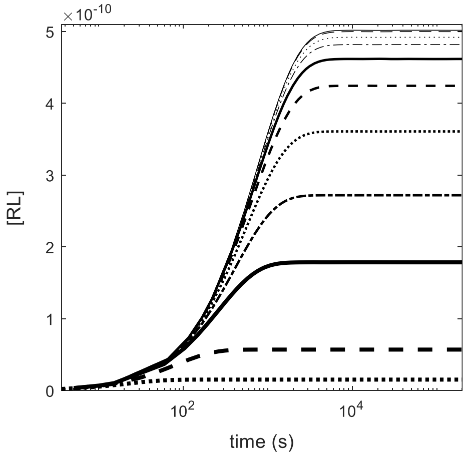


*Figure S7:* Semi-log plot of simulated of receptor-ligand association in the presence of increasing concentrations of inhibitor with $K_I$ = 4.9296E-10.

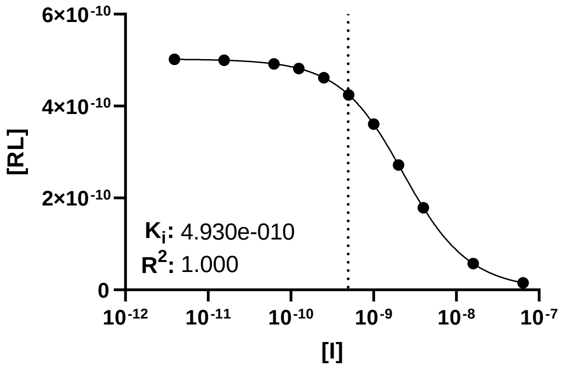


*Figure S8:* Curve-fitting of simulated competitive displacement titration data with equation S16. Data plotted semi-logx.

#### Section 2. Bivalent Binding and Ternary Complex Equilibria in Tight-Binding Conditions

In a homogenous solution, the association kinetics of mAbs with proteins is significantly complicated by formation of ternary complexes which occurs as some population of antibodies bind a second antigen. Often encountered in the development of immunoassays, the formation of these ternary complexes may be classified as examples of the 'high-dose hook effect' or 'antigen excess effect' that is sometimes referred to as a prozone/postzone phenomenon. The analytical intractability of the kinetics in these cases follows from the classical three-body problem, for which no general, closed form solution exists. One strategy to solve a three-body problem is reductive approximation, and we demonstrate that the analytical equations given by Morrison and Wang closely approximate the true solution-phase binding constants under certain conditions.

#### A, Mass-Balance Model of Tight-Binding Association of a Monoclonal Antibody

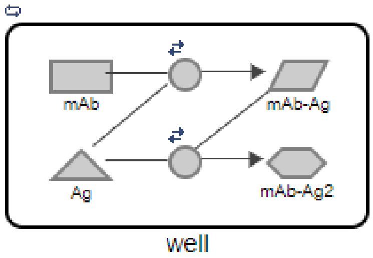


*Figure S9:* Graphical representation of a competitive displacement titration in which a receptor may be bound to a ligand or inhibitor.

$$\frac{d[AbAg]}{dt} = k_{on}[Ab][Ag] - k_{off}[AbAg] - k_f[AbAg][Ag] + k_r[AbAg2]$$

$$\frac{d[AbAg2]}{dt} = k_f[AbAg][Ag] - k_r[AbAg2]$$

$$\frac{d[Ab]}{dt} = k_{off}[AbAg] - k_{on}[Ab][Ag]$$

$$\frac{d[Ag]}{dt} = k_{off}[AbAg] - k_{on}[Ab][Ag] + k_r[AbAg2] - k_f[AbAg][Ag]$$

*Figure S10:* System of ODEs representing a competitive displacement titration. The interdependence of variables in this system implies that none can be fixed.

Association reactions of monoclonal antibodies and their antigens in solution may exhibit unpredictable behavior due to formation of ternary complexes. To model the effect of ternary complex formation on mAb-Ag association, MatLab Simbiology software was used to generate ODE-based models for the titration of mAb to a fixed concentration of antigen. A graphical model of the interaction was produced (Figure S9) and a system of ODEs was calculated for the model. (Figure S10) To elucidate the sources of error in applying the Morrison equation to this system, a simulation was performed in “tight binding” conditions at an antigen concentration exceeding the $K_D$. A numeric solver (ODE15s) within MatLab simulated a plot of the data over a range mAb concentrations. The formation of the binary mAb-Ag complex, as well as the total specific binding including the ternary complex are shown over time. (Figure S11) As may be observed by the hook-like inflections in the binding curve, ternary complex formation alters the equilibrium concentrations, irrespective of whether the analyte signal is the total bound mAb or the specific binary complex. We have observed similar inflections experimentally in the binding curves of trastuzumab labeled with AlexaFluor-488 and HER2-ECD protein labeled with Tb-cryptate. (Figure S12)

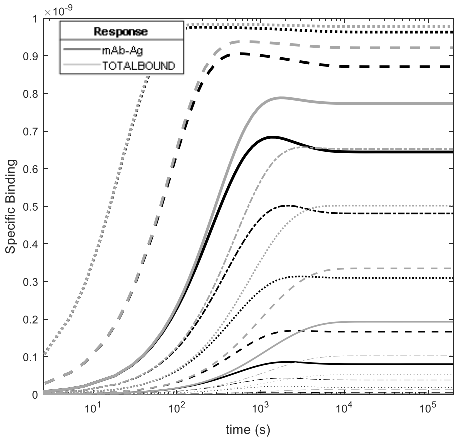


*Figure S11:* Semi-log plot of simulated bivalent mAb-antigen binding in the presence of increasing concentrations of labeled mAb. Equilibration of the mAb-Ag complex is illustrated in black, while the total of bound species including the ternary complex is illustrated in grey.

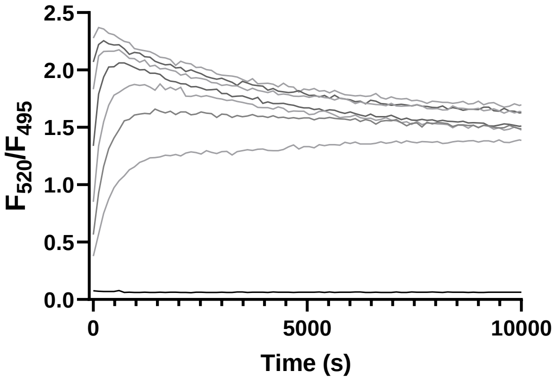


*Figure S12:* Association kinetics of trastuzumab-AF488 with Tb-labeled rHER2-ECD protein at 20ºC observed by TR-FRET on Synergy H1 instrument. Increasing concentrations of AF-488 labeled trastuzumab (0-16 nM) were the titrant to 1 nM preequilibrated HER2-anti-His Tb complex.

#### B. Analytical Intractability of Tight-Binding Association

Exact analytical equations are required to account for depletion of tight-binding ligands, yet no exact formula exists to describe the formation of ternary complexes in solution. Despite this limitation, mass-balance principles and reductive approximation offer significant insight. The observed rate constant of an association reaction between mAb and antigen in homogenous solution varies significantly with the concentration of antigen and deviates from the true value in tight-binding conditions. To model the effect of antigen concentration on apparent $K_D$ in this scenario, a series of simulations were performed in which a mAb was the titrant to an increasing series of fixed antigen concentrations. The apparent $K_D$ values were determined with Equation S10 and plotted against antigen concentration. (Figure S13) Important boundary conditions are illustrated by the plot, as the apparent $K_D$ derived from the Morrison equation deviates from its true value in ligand depletion regimes. These boundaries are also useful to delineate the 'prozone' and 'postzone' antigen concentration regimes. If the antigen concentration is less than the value of $K_D$, the apparent $K_D$ approximates the $K_D$ and the condition is prozone. As the antigen concentration approaches or exceeds the true $K_D$ value, the plots of apparent $K_D$ values exhibit non-linear increase, indicating postzone conditions among the titration series. Another boundary occurs as the antigen concentration approaches saturation and the assay condition is fully postzone. If $K_D$ is calculated as a function the total bound mAb, the plot reaches an inflection at which rapid decrease occurs due to depletion of free mAb. When $K_D$ is calculated as a function of the binary complex, the plot of apparent $K_D$ reaches an inflection point at which rapid increase occurs due to depletion of the analyte signal. This distinction may be relevant to spectroscopy if the FRET efficiencies of the donor species in binary and ternary complexes are not equivalent.

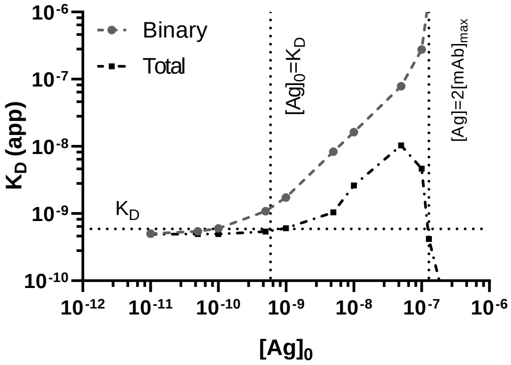


*Figure S13:* Log-log plot of apparent $K_D$ values obtained from a series of simulated mAb-Ag association reactions at increasing fixed concentrations of antigen. Activity of total bound antibody and specific mAb-Ag activity is used to derive apparent dissociation constants from equation S10.

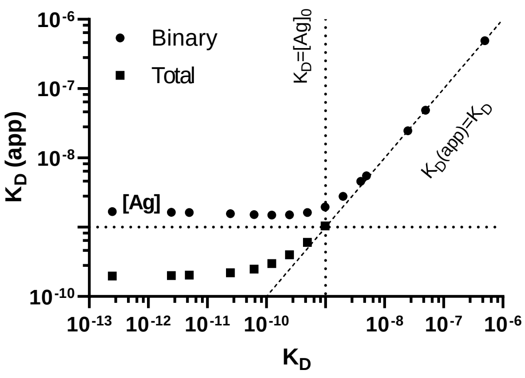


*Figure S14:* Log-log plot of apparent $K_D$ values obtained from a series of simulated mAb-Ag association reactions at increasing true $K_D$ values.

The utility of rate constants to distinguish prozone/postzone concentration regimes is illustrated clearly by comparing the apparent $K_D$ values of mAbs with a broad series of affinities. To model the effect of $K_D$ on the apparent $K_D$ of the binary mAb-Ag complex, simulations were performed in which a series of mAbs of increasing affinity were the titrant to a fixed 1 nM concentration of antigen. The $K_D$ values were determined with Equation S10 and plotted against the true affinity constant. (Figure S14) The plot shows that the apparent $K_D$ is equal to the true rate constant only as it exceeds the concentration of antigen. If the concentration of antigen is greater than the $K_D$, the apparent rate constant becomes tightly range-bound, even if the mAbs being evaluated differ in affinity by several orders of magnitude. This emphasizes the importance of the antigen concentration regime. When equivalence is the expected result, as is the case with the quality control of mAbs and ADCs, the potential for false positives is particularly tedious. If an experiment screens for high-affinity mAbs following affinity maturation techniques, the potential for false-negatives may cause the best candidates to be overlooked. We emphasize that development of immunological assays with mAbs should always be informed by the rate constant when it is available. Assay sensitivity may restrict the minimum concentration of reactants, particularly for miniaturized, high-throughput formats. As the value of $K_D$ often is not known *a priori*, the association kinetics of mAbs in solution should be evaluated with caution. The intractable coincidence of ligand depletion and ternary complex formation sufficiently accounts for the dominance of surface-based measurements in prior art. However, we further demonstrate that a competition assay format is not susceptible to these same type-I/type-II errors.

#### C. Mass-Balance Model of Competitive Displacement of Fab and mAb

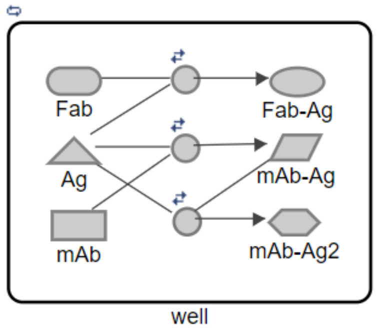


$$\frac{d[AbAg]}{dt} = k_{f1}[Ab][Ag] - k_{r1}[AbAg] + k_{r2}[AbAg2] - k_{f2}[Ab][AbAg]$$

$$\frac{d[AbAg2]}{dt} = k_{f2}[AbAg][Ag] - k_{r2}[AbAg2]$$

$$\frac{d[Ab]}{dt} = k_{r1}[AbAg] - k_{f1}[Ab][Ag]$$

$$\frac{d[FabAg]}{dt} = k_{on}[Fab][Ag] - k_{off}[FabAg]$$

$$\frac{d[Fab]}{dt} = k_{off}[FabAg] - k_{on}[Fab][Ag]$$

$$\frac{d[Ag]}{dt} = k_{off}[FabAg] - k_{on}[Fab][Ag] + k_{r1}[AbAg] - k_{f1}[Ab][Ag] + k_{r2}[AbAg2] - k_{f2}[Ag][AbAg]$$

*Figure S15:* Graphical representation of a competitive displacement titration in which a receptor may be bound to a ligand or inhibitor.

*Figure S16:* System of ODEs representing a competitive displacement titration for use in mass-balance simulation. The interdependence of variables in this system implies that none can be fixed.

When it is not possible to measure the binding affinity of a ligand directly, it is expedient to measure the displacement of a labeled ligand by the ligand of interest. To model the effect of ternary complex formation on competitive displacement, MatLab Simbiology software was used to generate ODE-based models for the titration of increasing concentrations of mAb to a fixed concentration of its fragment of antigen binding. A graphical model of the interaction was produced (Figure S15) and a system of ODEs was calculated for the model. (Figure S16) A numeric solver (ODE15s) simulated a plot of the data corresponding to the equilibrium concentration of the Fab-Ag complex at each concentration of mAb with the same rate-constants used in previous experiments, and the apparent $K_I$ of the mAb was calculated with Equation S16. The simulated titration was repeated across a series of antigen concentrations. (Figure S17) and across a series of $K_I$ values at a fixed 1 nM concentration of antigen. (Figure S18)

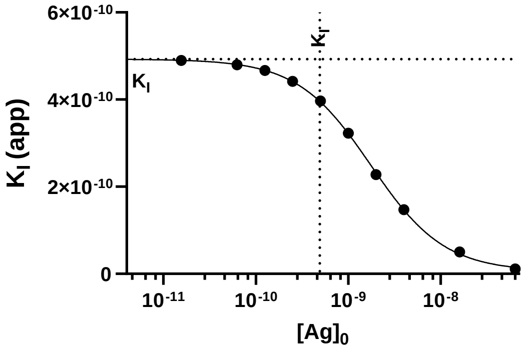


*Figure S17:* Log-log plot of apparent $K_I$ values obtained from a series of simulated competitive displacement titration of Fab by mAb at increasing fixed concentrations of antigen.

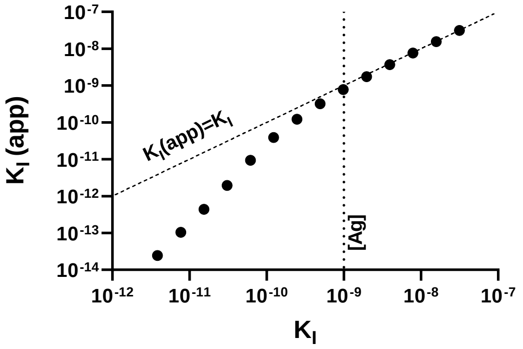


*Figure S18:* Log-log plot of apparent $K_I$ values obtained from a series of simulated competitive displacement titration of Fab by mAb at increasing true $K_I$ values.

A comparison of the plots outlined in Figure S14 and Figure S18 demonstrated that the competition assay format is superior for measurement of tight-binding affinities in homogenous solution. These plots show that if the antigen concentration exceeds the equilibrium constant, the apparent constant will deviate from the true value. However, the slope of deviation from the line of identity in Figure S18 (competition format) is significantly more acute than that of Figure S14 (direct measurement) with the same kinetic parameters. The error induced by tight-binding is mitigated, as the effective binding affinity of the mAb is reduced by competition. Importantly, the competition assay format demonstrates no ambiguity in the rank-order of the analytes, even under extreme conditions of depletion. Unlike direct the measurements of mAb $K_D$ illustrated in Figure S14, there is no false equivalence among apparent $K_I$ values in tight-binding conditions. Therefore, measurement of the $K_I$ of a mAb against its own Fab appears to eliminate the associated risk of type-I/type-II errors. Accordingly, competitive displacement titration was selected as the format of the assay.

#### Section 3. Competitive Displacement TR-FRET Assay for Evaluating Antibody Affinity

##### A. Graphical Overview

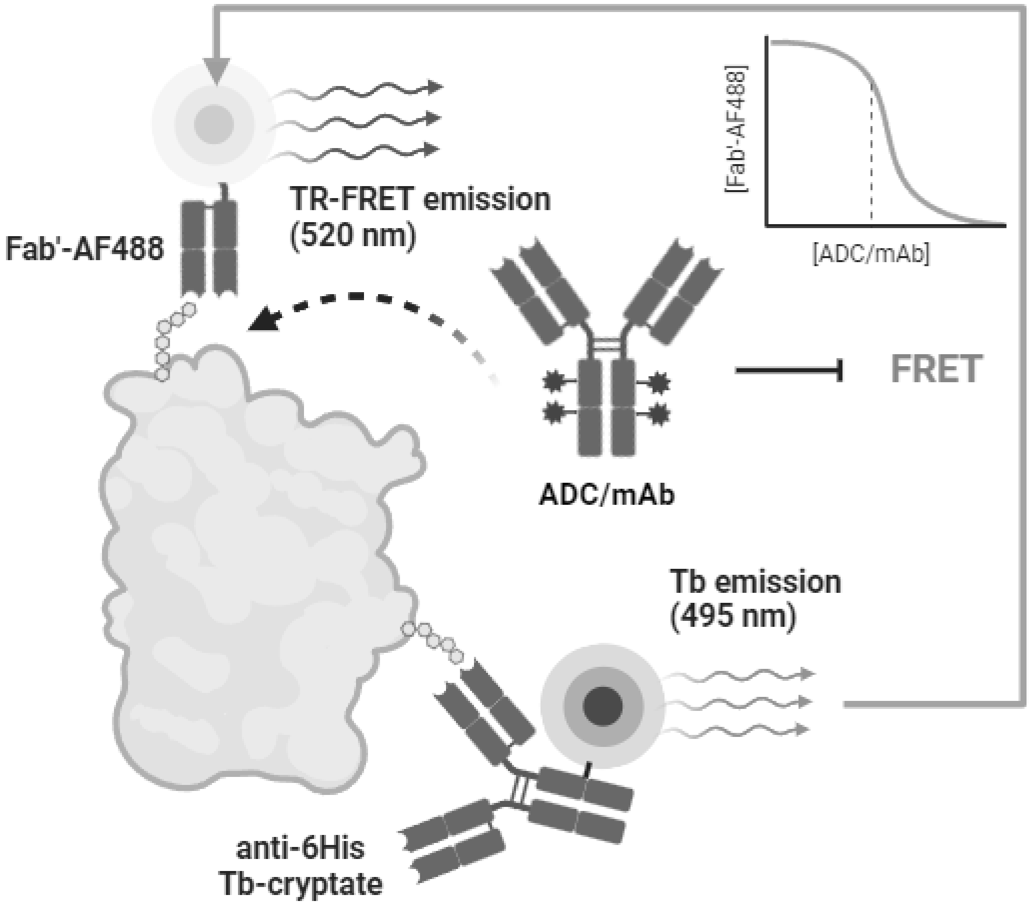


**Scheme S1:** Inhibition of observed FRET interaction between a Fab'-AF488 tracer and anti-6His Tb-cryptate labeled protein by an ADC or mAb[25]

#### B. Synthesis of Fab'-Alexafluor488 TR-FRET Tracer

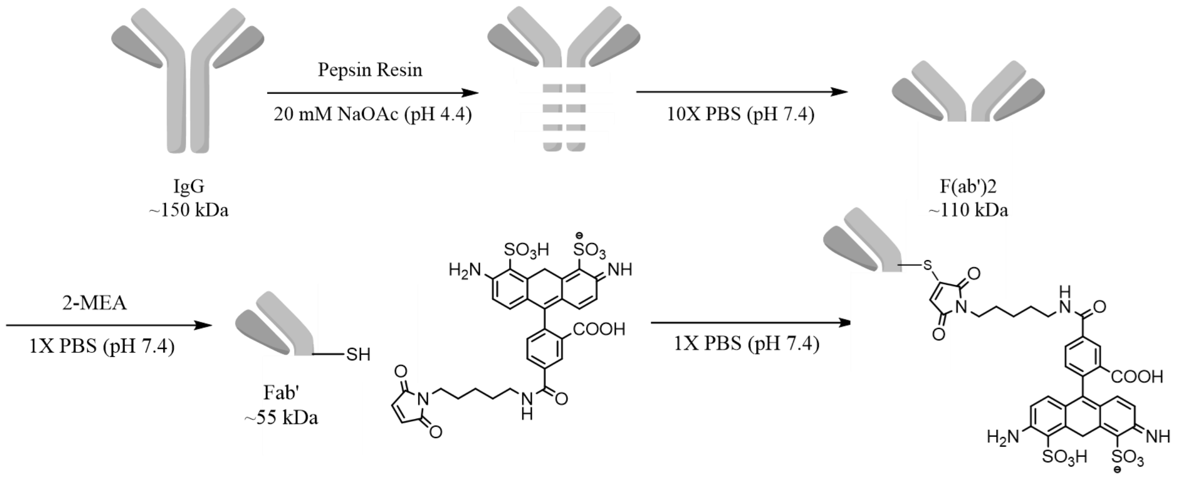


Scheme S2: Synthesis of trastuzumab Fab'-Alexafluor 488

An empty 1 mL spin column was loaded with 250 uL of immobilized pepsin/agarose solution, and centrifuged at 5000xg for 1 min. The compacted resin was washed twice and resuspended with 0.5 mL 20 mM NaOAc at pH 4.4. Trastuzumab (Herceptin®, Genentech) at 21 mg/mL was buffer exchanged into 20 mM NaOAc at pH 4.4 in a 0.5 mL Zeba™ spin column and assayed by BCA. An aliquot of 2 mg purified trastuzumab was added to 20 mM NaOAc at pH 4.4 to a total volume of 0.5 mL and combined with the suspension of immobilized pepsin in the spin column, which was capped and incubated at 37ºC with shaking for 2h. The column was centrifuged at 5000xg for 1 min and washed twice with 0.5 mL 10X PBS. The supernatant and washes containing trastuzumab F(ab')2 fragments were combined, and buffer exchanged into 0.5 mL 10 mM EDTA in 1X PBS with a 10k MWCO Amicon Ultra centrifugal filter. A solution of 2 mM 2-mercaptoethylamine was prepared in 1X PBS. An aliquot of 2-MEA (2 eq) was added to the crude F(ab')2 and the solution was incubated at 37ºC for 90 min. The solution was cooled, adjusted to 15% DMSO, and an excess (4 eq) of AlexaFluor® 488 maleimide was added, followed by incubation at room temperature for 2h. The reaction was buffer exchanged into 1X PBS and purified by FPLC on a Superdex® 200 (30 cm/10 mm) column. Those fractions with absorbance at 488 nm were characterized by gel electrophoresis. (Figure S19) Fractions corresponding to the expected mass ~45 kDa were isolated and concentrated. The concentration of Fab'-AF488 was determined by absorbance at 280 nm (ε=68,590 $M^{-1}cm.$)[1] Aliquots were frozen in liquid nitrogen and stored at -80ºC for further experiments.

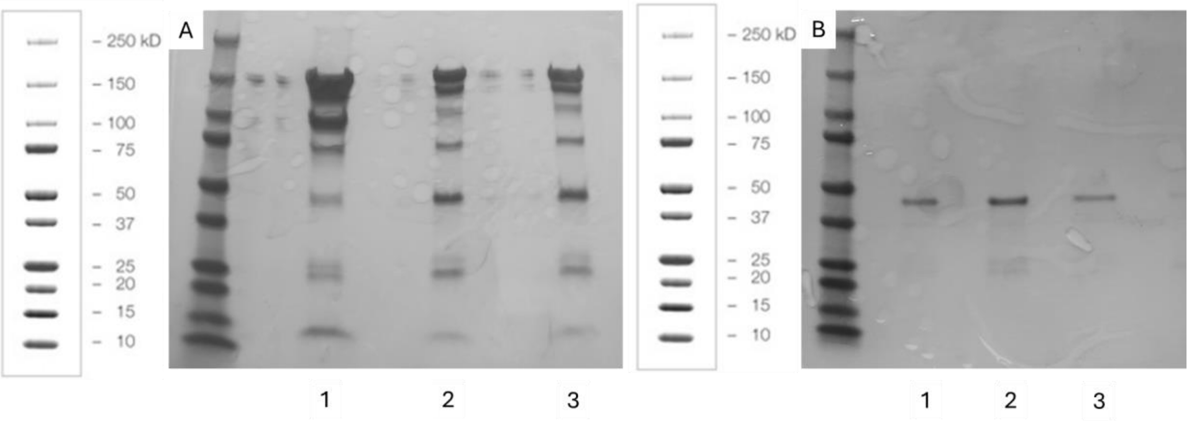

Figure S19: (A) PAGE of (1) Pepsin digestion of trastuzumab (2) 2-MEA reduction of crude digestion (3) Labeling of crude digestion (B) PAGE of FPLC fractions (1,2,3) containing Fab'-AF488

#### C. Evaluating Drug-to-Antibody Ratio / Degree of Labeling by UV Spectroscopy

The average drug-to-antibody ratio (DAR), or degree of labeling (DOL) was calculated for each species from UV measurements at the wavelength of maximum absorption for each species with the protocol given by Chen.[24] The molar absorptivity values for conjugates of vc-MMAE, smcc-DM1, and Alexafluor 488 were taken from literature values reported in Table S1 and used to determine the relative concentrations of antibody and label. Briefly, the individual contributions of the antibody and drug to measured absorbance were determined by simultaneously solving the system of Equation S17 and Equation S18.

$$A_{280} = \left(\varepsilon_{drug}^{280} c_{drug} + \varepsilon_{mAb}^{280} c_{mAb}\right) l \quad \text{(S17)}$$

$$A_{\lambda(D)} = \left(\varepsilon_{drug}^{\lambda(D)} c_{drug} + \varepsilon_{mAb}^{\lambda(D)} c_{mAb}\right) l \quad \text{(S18)}$$

$$c_{mAb} = \left(A_{280}\varepsilon_{drug}^{\lambda(D)} - A_{\lambda(D)}\varepsilon_{drug}^{280}\right) / \left(\left(\varepsilon_{mAb}^{280}\varepsilon_{drug}^{\lambda(D)} - \varepsilon_{mAb}^{\lambda(D)}\varepsilon_{drug}^{280}\right) l\right) \quad \text{(S19)}$$

$$c_{drug} = \left(A_{280}\varepsilon_{mAb}^{\lambda(D)} - A_{\lambda(D)}\varepsilon_{mAb}^{280}\right) / \left(\left(\varepsilon_{drug}^{280}\varepsilon_{mAb}^{\lambda(D)} - \varepsilon_{drug}^{\lambda(D)}\varepsilon_{mAb}^{280}\right) l\right)$$

$$DAR = c_{drug}/c_{mAb} \quad \text{(S20)}$$

**Table S1: Molar Absorptivity of Trastuzumab and Labels**

| | ε248 | ε252 | ε280 | ε488 |
|---|---|---|---|---|
| Tmab[2] | 7.75E+04 | 7.66E+04 | 2.25E+05 | 0.00E+00 |
| vc-MMAE[2] | 1.50E+03 | - | 1.59E+04 | - |
| smcc-DM1[3] | - | 2.68E+04 | 5.70E+03 | - |
| AF-488[4] | - | - | 0.00E+00 | 7.30E+04 |

#### D. General Protocol for Conducting Homogenous TR-FRET Assay

The assay was conducted in a 384-well white microplate at a volume of 20 μL/well. All pipetting was performed by the Formulatrix Mantis® liquid dispensing robot. The buffer consisted of equal parts reaction buffer with 50 mM Tris at pH 7.5, 1 mM TCEP, 0.1% Triton X-100, and 0.01% BSA in dH2O and storage buffer with 1X PBS at pH 7.4. Series dilution of the analytes was performed in a 96-well plate according to the number of test wells. Dilution schemes were optimized around a median concentration on the order of the anticipated rate constant. The dose-response curve of each analyte was evaluated in triplicate alongside a negative control lacking only the antigen. In each well, the analyte and tracer were pipetted first, followed by the Tb-labeled antigen complex or control Tb fluorophore. Reagents were prepared from stock at 4X working concentrations and pipetted into each 20 μL well in 5 μL volumes. The volume in each well was normalized to contain 10 μL reaction buffer and 10 μL 1X PBS. To prepare the Tb-labeled receptor solution for all test wells, a 2.5 mL solution containing 2 nM HER2-ECD and 2.4 nM anti-His Tb-cryptate in 1X PBS was incubated at 4ºC for 72 h. Tracer-antigen association was measured in 3x12 test wells, each containing 5 uL of tracer at increasing concentrations and 0.5 nM Tb-labeled receptor. A 2.5 mL solution of tracer was prepared by diluting Fab'-AF488 to an appropriate concentration in reaction buffer. The affinity of each analyte was evaluated in competition with this fixed concentration of tracer, with 3x12 test wells containing 5 μL analyte at increasing concentrations, 5 μL tracer solution, and 0.5 nM Tb-labeled antigen. The microplate was sealed with film and incubated at 4ºC for the duration of the experiment. After the assay was equilibrated, or at appropriate intervals, the donor (495 nm) and acceptor (520 nm) emissions for each well were collected with excitation (340 nm) on a Biotek Synergy H1 microplate reader equipped with a LanthaScreen-Tb module. (EX:340/30, EM:495/10, EM:520/25).

#### E. Curve Fitting

The ratio of 520 and 495 nm emissions observed in each well was calculated and the non-specific background subtracted to prepare data for curve fitting. The FRET ratio for each series was normalized and plotted against analyte concentration. Curve fitting was performed in GraphPad Prism 9. The kinetic parameters of the tracer were determined by fitting the dose-response curve to Equation 1 where x is the concentration of Fab'-AF488 and y is the normalized specific binding estimated by the FRET ratio.

$$y = ([\mathrm{Ag}]_0 + \mathrm{x} + \mathrm{K_D}) - \sqrt{([\mathrm{Ag}]_0 + \mathrm{x} + \mathrm{K_D})^2 - 4[\mathrm{Ag}]_0\mathrm{x}}\,)/2[\mathrm{Ag}]_0 \quad \text{(Equation 1)}$$

The best fit parameters of the Fab $K_D$ and antigen concentration were adopted to fit the dose-response curves for each mAb analyte using Equation 2, where x is the concentration of mAb, and y is the normalized specific binding estimated by the FRET ratio. Apparent $K_I$ values were reported from the best-fit parameters of the curve. In cases that equilibration was monitored, $K_I$ values obtained in periodic measurements of each analyte were plotted against the precise time reported by the instrument. The plots were fitted with a one-phase decay model, and the plateaus were reported as the apparent KI values with asymmetric 95% confidence.

$$y = \frac{[FAb]_0\left\{2\sqrt{(a^2 - 3b)} * \cos\left(\frac{\theta}{3}\right) - a\right\}}{3K_D + \left\{2\sqrt{(a^2 - 3b)} * \cos\left(\frac{\theta}{3}\right) - a\right\}}/[Ag]_0 \quad \text{(Equation 2)}$$

where:

$a = K_D + K_I + [Fab] + x - [Ag]$

$b = K_I([Fab] - [Ag]) + K_D(x - [Ag]) + K_D K_I$

$c = -K_D K_I [Ag]$

$\theta = \arccos(-2a^3 + 9ab - 27c)/(2\sqrt{(a^2 - 3b)^3})$

### Section 4. Evaluation of the Apparent Affinity of Trastuzumab

#### A. Effect of Assay HER2-ECD Concentration in Apparent $K_I$ Evaluation

A series of competition assays were performed between 2 nM Fab'-AF488 and trastuzumab over a range of HER2-ECD antigen concentrations from 250 pM-4 nM. Following the general protocol in Supplementary Section 2E/F, the assays were equilibrated at 4ºC for 18 hours prior to collecting the data. The dose-response curves and fit are shown in Figure S20. Fortunately, we do not observe variation in the apparent $K_I$ of trastuzumab in the given conditions at HER2-ECD concentrations up to 1 nM, and this set of experiments demonstrates excellent precision with apparent $K_I$ value 10.80 pM ± 0.03 pM. In contrast, the observed $IC_{50}$ values increase with antigen concentration over this range. At the antigen concentrations of 2 nM and 4 nM, the expected decrease is apparent in $K_I$ values of 9.1 pM and 9.0 pM, respectively. These experiments validate the use of antigen concentrations of 250 pM-1 nM in the given assay condition. Further experiments were conducted at an HER2-ECD concentration of 500 pM.

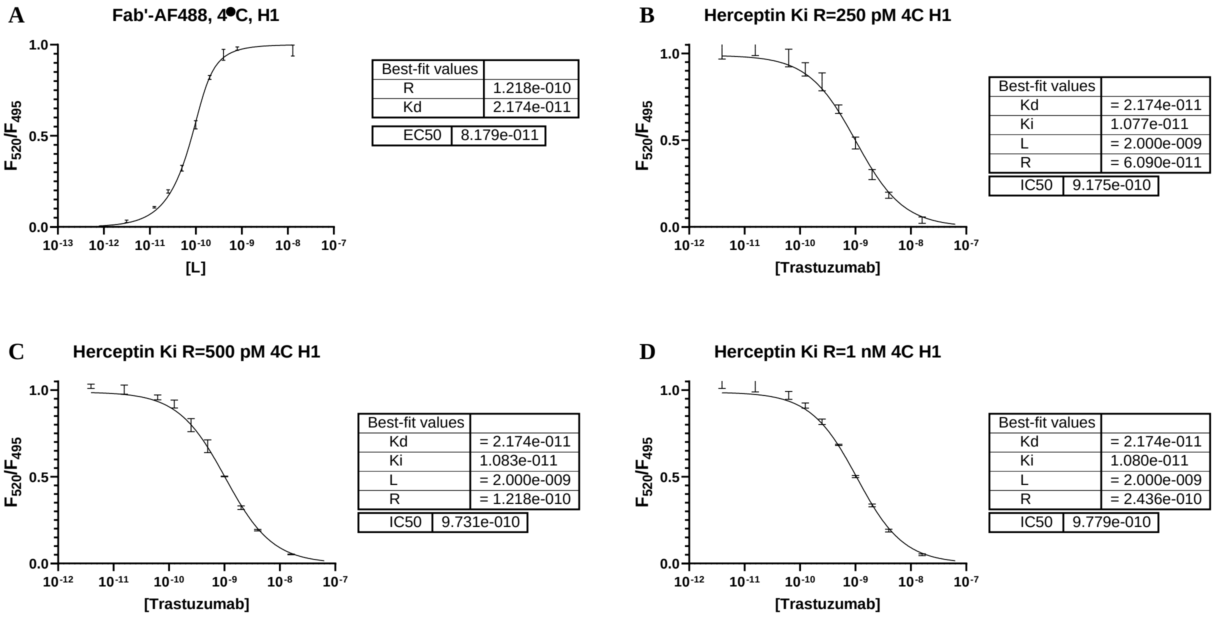

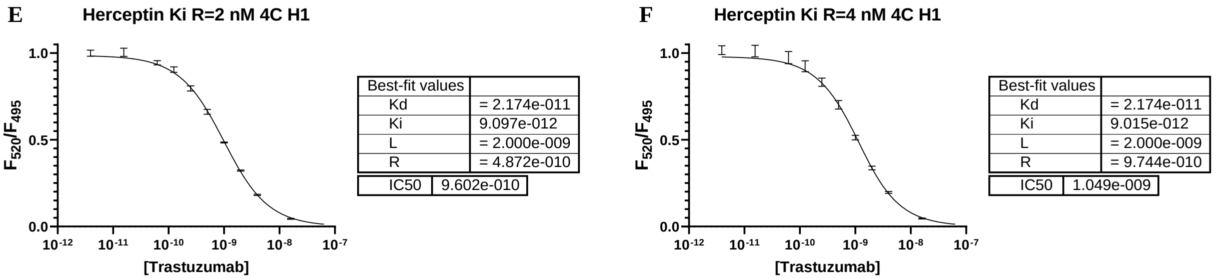


*Figure S20*: Effect of HER2-ECD Concentration in Apparent $K_I$ Evaluation. (A) Curve-fitting of Fab'-AF488 tracer. (B-F) Curve fitting of trastuzumab apparent KI at receptor concentrations 250 pM-1 nM.

#### B. Effect of Assay Fab Concentration on Apparent $K_I$

A series of competition assays were conducted between trastuzumab and 500 pM HER2-ECD antigen over a range of Fab'-AF488 concentrations from 2 nM-32 nM. Following Supplementary Section 2E, the assays were equilibrated for 40 h at 4ºC and the data collected at intervals during equilibration. Curve-fitting was performed according to the Supplementary Section 2F, and the $K_I$ values were fit by a one-phase decay model to give the reported apparent $K_I$. The one-phase decay model is shown in Figure S23. The dose-response curves and fits are shown in Figure S24-S27. The apparent $K_I$ of trastuzumab in the given conditions at Fab'-AF488 concentrations of 2 nM-32 nM decreases in linear fashion from 7.3 pM-4.4 pM. (Figure S22) Among three observations, linear regression gives $R^2$ of 1.000. These experiments likely illustrate photophysical interference. The linear relationship may permit an estimate of the scale of the error in the given assay condition by comparing the values to the y-intercept.

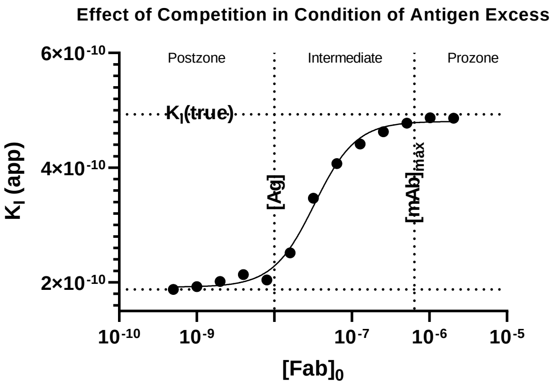


*Figure S21*: Predicted trend of apparent $K_I$ values obtained from numerical simulation with MatLab Simbiology of increasing Fab competitor in condition of antigen excess.

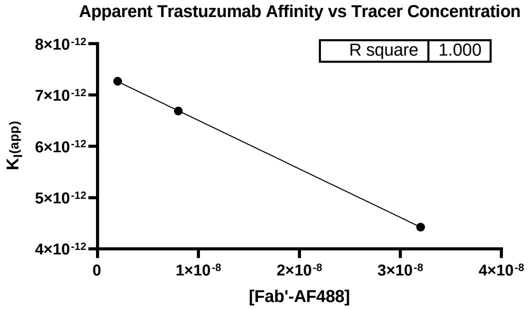


*Figure S22*: Linear trend in apparent $K_I$ values in assay experiments with increasing concentrations of Fab'-AF488.

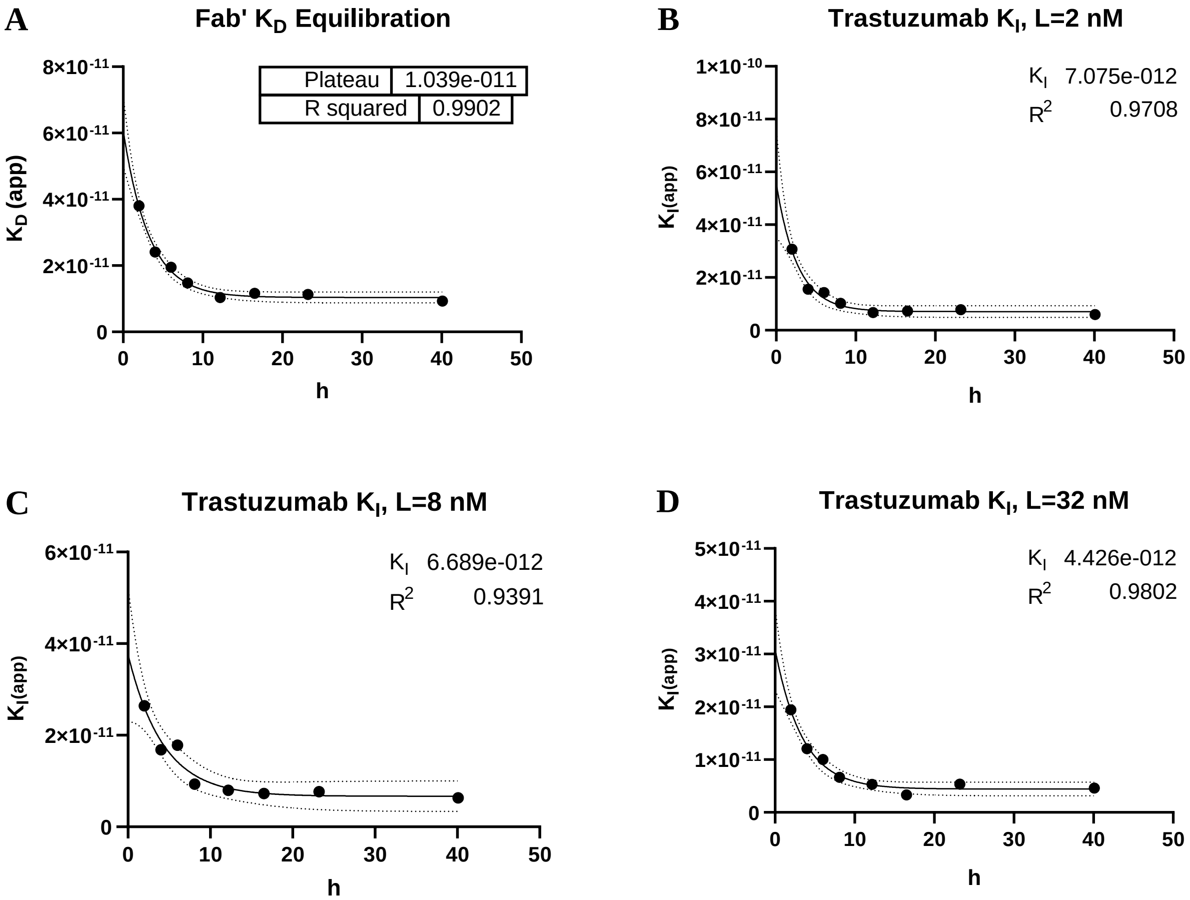


*Figure S23*: Effect of Tracer Concentration in Apparent $K_I$ Evaluation. (A) Curve-fitting of Fab'-AF488 tracer equilibration $K_I$ observations by one-phase decay. (B-D) Curve fitting of trastuzumab equilibration $K_I$ observations at tracer concentrations 2 nM-32 nM by one-phase decay.

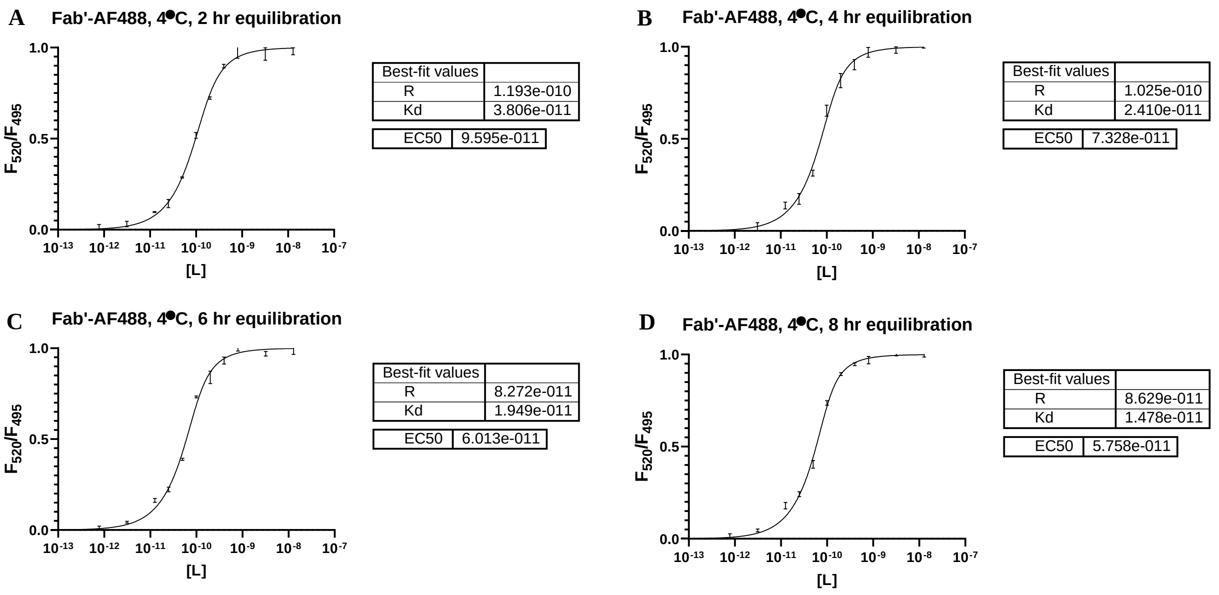

**E** **Fab'-AF488, 4●C, 12 hr equilibration**

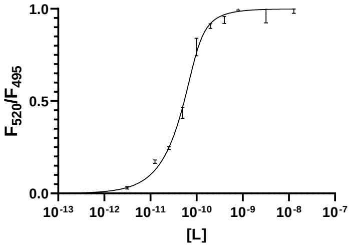


| Best-fit values | |
|---|---|
| R | 8.719e-011 |
| Kd | 1.039e-011 |

| | |
|---|---|
| EC50 | 5.119e-011 |

**F** **Fab'-AF488, 4●C, 16 hr equilibration**

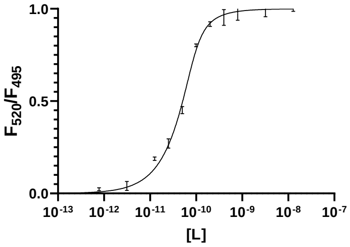


| Best-fit values | |
|---|---|
| R | 1.285e-016 |
| Kd | 1.161e-011 |

| | |
|---|---|
| EC50 | 1.276e-011 |

**G** **Fab'-AF488, 4●C, 23 hr equilibration**

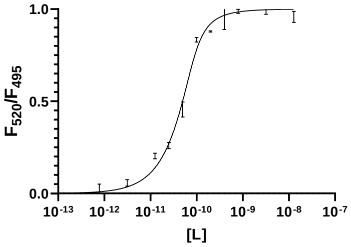


| Best-fit values | |
|---|---|
| R | 7.699e-011 |
| Kd | 1.128e-011 |

| | |
|---|---|
| EC50 | 4.867e-011 |

**H** **Fab'-AF488, 4●C, 40 hr equilibration**

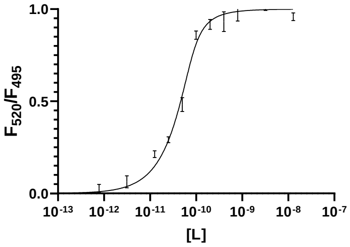


| Best-fit values | |
|---|---|
| R | 7.372e-011 |
| Kd | 9.315e-012 |

| | |
|---|---|
| EC50 | 4.468e-011 |

*Figure S24*: Effect of Tracer Concentration in Apparent $K_I$ Evaluation. Dose response curves and fitting of Fab'-AF488 association with HER2-ECD at time points (A) 2h (B) 4h (C) 6h (D) 8h (E) 12h (F) 16h (G) 23h (H) 40 h

**A** **Herceptin Ki, 4●C, 4 hr equilibration**

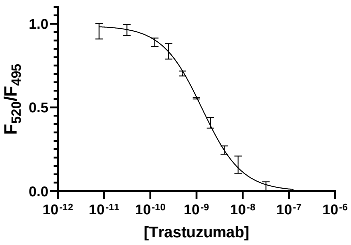


| Best-fit values | |
|---|---|
| Kd | = 2.410e-011 |
| Ki | 1.550e-011 |
| L | = 2.000e-009 |
| R | = 1.025e-010 |

| | |
|---|---|
| IC50 | 1.420e-009 |

**B** **Herceptin Ki, 4●C, 6 hr equilibration**

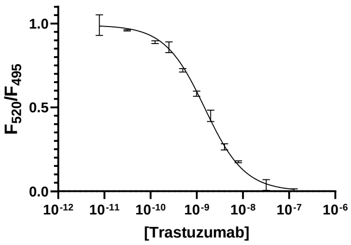


| Best-fit values | |
|---|---|
| Kd | = 1.949e-011 |
| Ki | 1.433e-011 |
| L | = 2.000e-009 |
| R | = 8.272e-011 |

| | |
|---|---|
| IC50 | 1.558e-009 |

**C** **Herceptin Ki, 4●C, 8 hr equilibration**

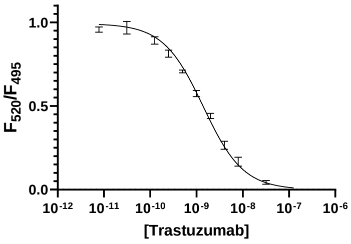


| Best-fit values | |
|---|---|
| Kd | = 1.478e-011 |
| Ki | 1.020e-011 |
| L | = 2.000e-009 |
| R | = 8.629e-011 |

| | |
|---|---|
| IC50 | 1.544e-009 |

**D** **Herceptin Ki, 4●C, 12 hr equilibration**

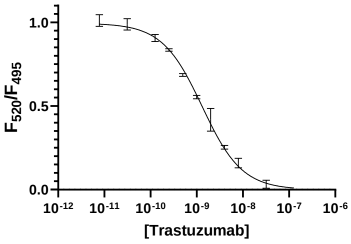


| Best-fit values | |
|---|---|
| Kd | = 1.039e-011 |
| Ki | 6.696e-012 |
| L | = 2.000e-009 |
| R | = 8.719e-011 |

| | |
|---|---|
| IC50 | 1.273e-009 |

**E** **Herceptin Ki, 4●C, 16 hr equilibration**

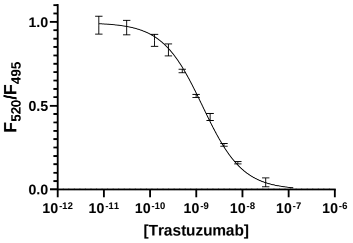


| Best-fit values | |
|---|---|
| Kd | = 1.063e-011 |
| Ki | 7.255e-012 |
| L | = 2.000e-009 |
| R | 9.549e-012 |

| | |
|---|---|
| IC50 | 1.450e-009 |

**F** **Herceptin Ki, 4●C, 23 hr equilibration**

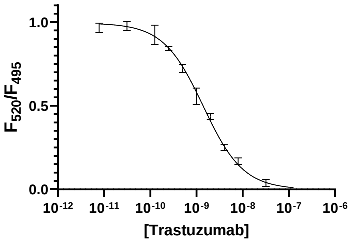


| Best-fit values | |
|---|---|
| Kd | = 1.128e-011 |
| Ki | 7.807e-012 |
| L | = 2.000e-009 |
| R | = 7.699e-011 |

| | |
|---|---|
| IC50 | 1.449e-009 |

**G** **Herceptin Ki, 4●C, 40 hr equilibration**

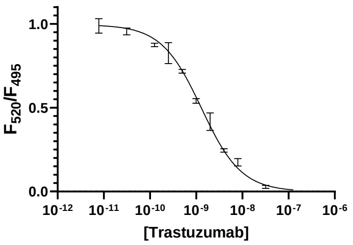


| Best-fit values | |
|---|---|
| Kd | = 9.315e-012 |
| Ki | 5.960e-012 |
| L | = 2.000e-009 |
| R | = 7.372e-011 |
| IC50 | 1.393e-009 |

**Herceptin Ki, 4●C, 2 hr equilibration**

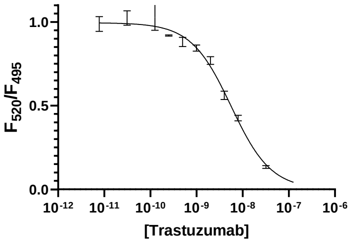


| Best-fit values | |
|---|---|
| Kd | = 3.806e-011 |
| Ki | 2.642e-011 |
| L | = 8.000e-009 |
| R | = 1.193e-010 |
| IC50 | 6.515e-009 |

*Figure S25*: Effect of Tracer Concentration in Apparent $K_I$ Evaluation. Dose response curves and fitting of 2 nM Fab'-AF488 disassociation from HER2-ECD with increasing trastuzumab concentration at time points (A) 2h (B) 4h (C) 6h (D) 8h (E) 12h (F) 16h (G) 23h (H) 40 h

**A** **Herceptin Ki, 4●C, 2 hr equilibration**

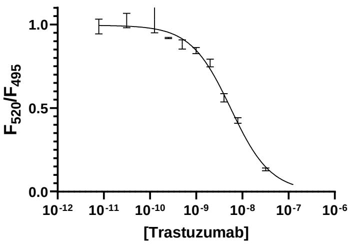


| Best-fit values | |
|---|---|
| Kd | = 3.806e-011 |
| Ki | 2.642e-011 |
| L | = 8.000e-009 |
| R | = 1.193e-010 |
| IC50 | 6.515e-009 |

**B** **Herceptin Ki, 4●C, 4 hr equilibration**

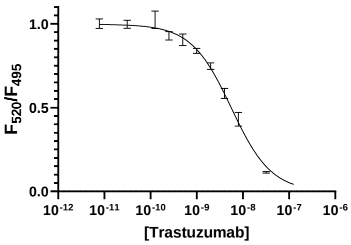


| Best-fit values | |
|---|---|
| Kd | = 2.410e-011 |
| Ki | 1.680e-011 |
| L | = 8.000e-009 |
| R | = 1.025e-010 |
| IC50 | 6.625e-009 |

**C** **Herceptin Ki, 4●C, 6 hr equilibration**

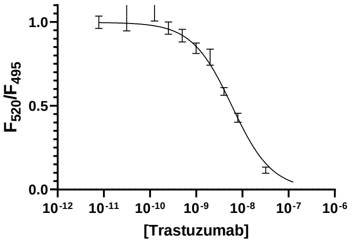


| Best-fit values | |
|---|---|
| Kd | = 2.410e-011 |
| Ki | 1.782e-011 |
| L | = 8.000e-009 |
| R | = 8.272e-011 |
| IC50 | 6.268e-009 |

**D** **Herceptin Ki, 4●C, 8 hr equilibration**

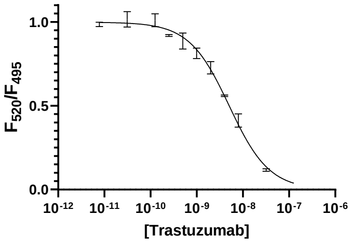


| Best-fit values | |
|---|---|
| Kd | = 1.478e-011 |
| Ki | 9.307e-012 |
| L | = 8.000e-009 |
| R | = 8.629e-011 |
| IC50 | 6.159e-009 |

**E** **Herceptin Ki, 4●C, 12 hr equilibration**

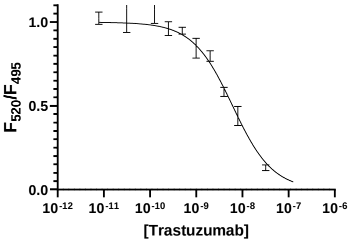


| Best-fit values | |
|---|---|
| Kd | = 1.039e-011 |
| Ki | 7.969e-012 |
| L | = 8.000e-009 |
| R | = 8.719e-011 |
| IC50 | 6.255e-009 |

**F** **Herceptin Ki, 4●C, 16 hr equilibration**

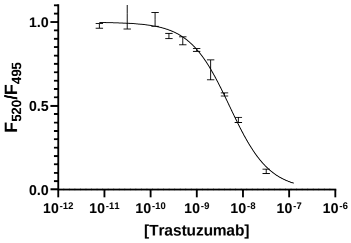


| Best-fit values | |
|---|---|
| Kd | = 1.161e-011 |
| Ki | 7.299e-012 |
| L | = 8.000e-009 |
| R | 4.343e-010 |
| IC50 | 6.125e-009 |

**G** **Herceptin Ki, 4●C, 23 hr equilibration**

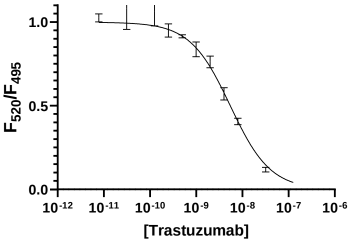


| Best-fit values | |
|---|---|
| Kd | = 1.128e-011 |
| Ki | 7.694e-012 |
| L | = 8.000e-009 |
| R | = 7.699e-011 |
| IC50 | 5.674e-009 |

**H** **Herceptin Ki, 4●C, 40 hr equilibration**

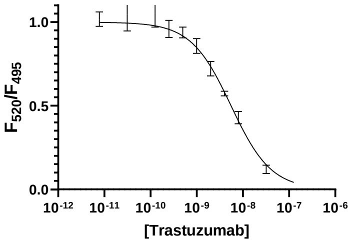


| Best-fit values | |
|---|---|
| Kd | = 9.135e-012 |
| Ki | 6.308e-012 |
| L | = 8.000e-009 |
| R | = 7.372e-011 |
| IC50 | 5.819e-009 |

*Figure S26*: Effect of Tracer Concentration in Apparent $K_I$ Evaluation. Dose response curves and fitting of 8 nM Fab'-AF488 disassociation from HER2-ECD with increasing trastuzumab concentration at time points (A)

2h (B) 4h (C) 6h (D) 8h (E) 12h (F) 16h (G) 23h (H) 40 h

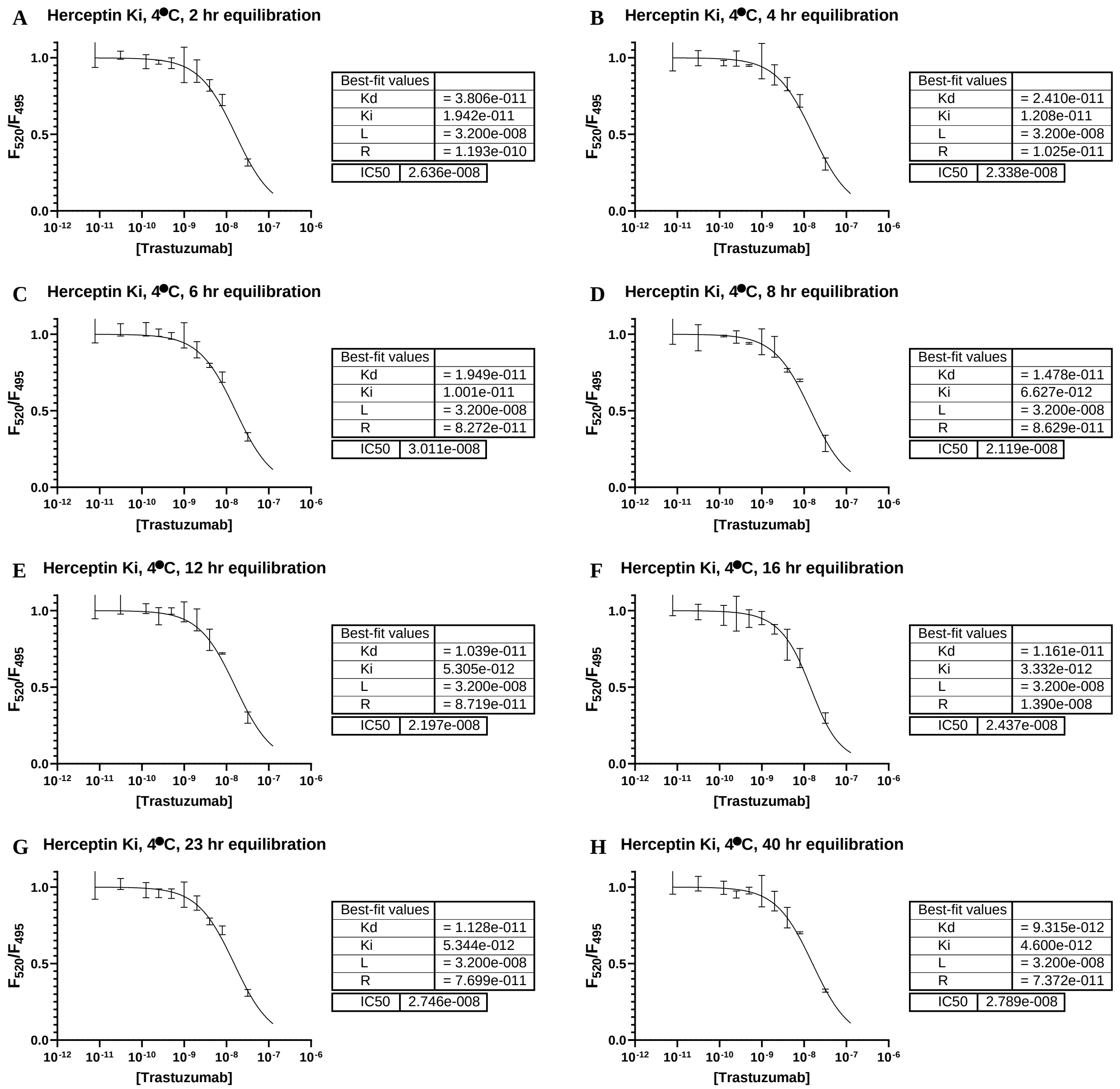


*Figure S27*: Effect of Tracer Concentration in Apparent $K_I$ Evaluation. Dose response curves and fitting of 32 nM Fab'-AF488 disassociation from HER2-ECD with increasing trastuzumab concentration at time points (A) 2h (B) 4h (C) 6h (D) 8h (E) 12h (F) 16h (G) 23h (H) 40 h

### Section 5. Evaluation of Conditions Relevant to Production of Antibody-Drug Conjugates

#### A. Effect of Solvent Exposure on the Binding Affinity of Trastuzumab

A 1 mg aliquot of trastuzumab in 1X PBS was added into microcentrifuge tubes containing 1X PBS and 0%, 10%, 25%, or 50% organic (DMF/DMSO) in a final volume of 200 µL. The samples were placed on a shaker and incubated at room temperature for 2h. Each sample was then buffer exchanged into 1X PBS on a 10K MWCO 0.5 mL Amicon® centrifugal filter and purified by gel filtration. Concentrations were determined by UV absorbance at 280 nm. Each sample was frozen in liquid N2 and stored at -80ºC for further analysis. Following the protocol in Supplementary Section 2E/F, competition assays were equilibrated for 24 h at 4ºC with test wells containing 2 nM Fab'-AF488 and 0.5 nM HER2-ECD antigen. Curve-fitting was performed according to the general protocol in Supplementary Section 2F. The results are summarized in Figure S28. The dose-response curves and fits are shown in Figure S29.

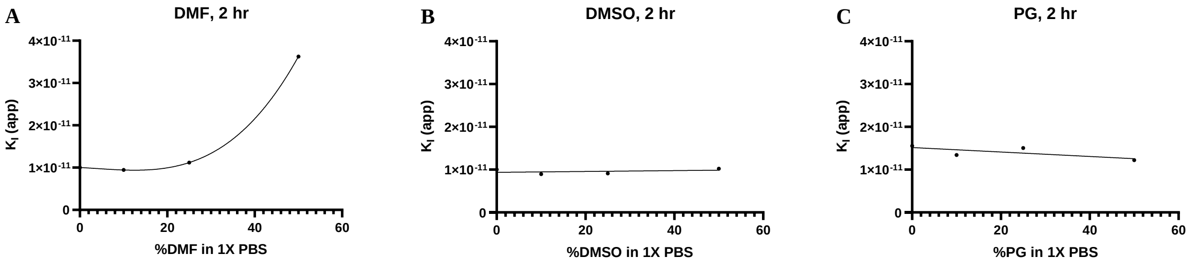


*Figure S28*: Effect of 2 h exposure to organic solvent on the apparent KI of trastuzumab in 1X PBS at room temperature. (A) Dimethylformamide (B) Dimethyl sulfoxide (C) Propylene Glycol

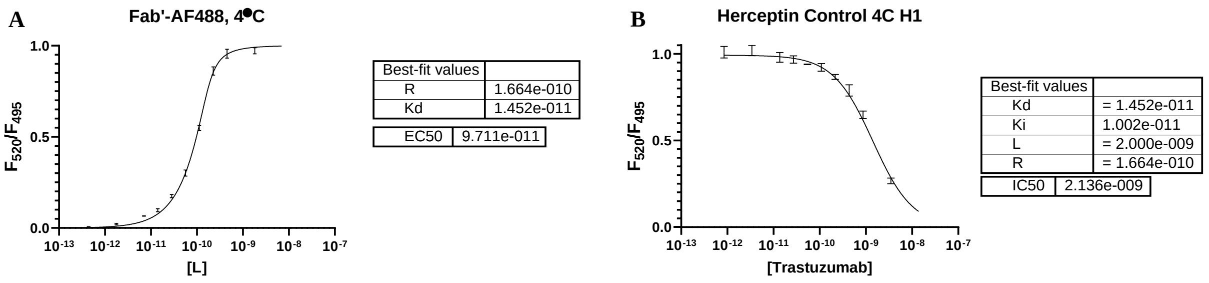

**C** Herceptin DMF 10%

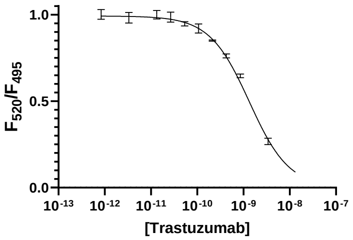


| Best-fit values | |
|---|---|
| Kd | = 1.452e-011 |
| Ki | 9.435e-012 |
| L | = 2.000e-009 |
| R | = 1.664e-010 |
| IC50 | 2.095e-009 |

**D** Herceptin 25% DMF

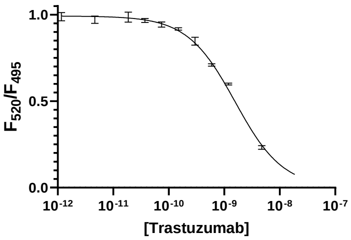


| Best-fit values | |
|---|---|
| Kd | = 1.452e-011 |
| Ki | 1.118e-011 |
| L | = 2.000e-009 |
| R | = 1.664e-010 |
| IC50 | 2.179e-009 |

**E** Herceptin 50% DMF 4C H1

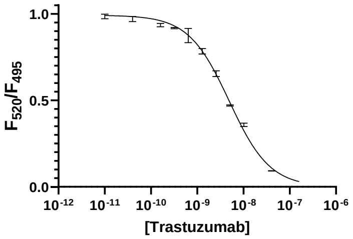


| Best-fit values | |
|---|---|
| Kd | = 1.452e-011 |
| Ki | 3.626e-011 |
| L | = 2.000e-009 |
| R | = 1.664e-010 |
| IC50 | 5.741e-009 |

**F** Herceptin 10% DMSO 4C H1

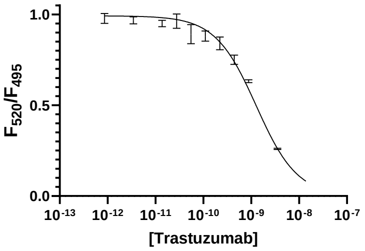


| Best-fit values | |
|---|---|
| Kd | = 1.452e-011 |
| Ki | 8.964e-012 |
| L | = 2.000e-009 |
| R | = 1.664e-010 |
| IC50 | 2.277e-009 |

**G** Herceptin 25% DMSO 4C H1

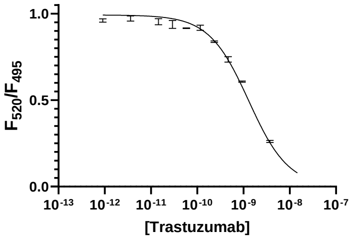


| Best-fit values | |
|---|---|
| Kd | = 1.452e-011 |
| Ki | 9.121e-012 |
| L | = 2.000e-009 |
| R | = 1.664e-010 |
| IC50 | 2.075e-009 |

**H** Herceptin 50% DMSO 4C H1

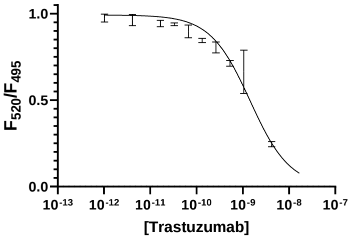


| Best-fit values | |
|---|---|
| Kd | = 1.452e-011 |
| Ki | 1.022e-011 |
| L | = 2.000e-009 |
| R | = 1.664e-010 |
| IC50 | 3.259e-009 |

**I** Fab'-AF488, 4●C, H1

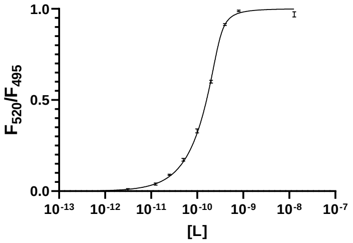


| Best-fit values | |
|---|---|
| R | 2.946e-010 |
| Kd | 1.248e-011 |
| EC50 | 1.512e-010 |

**J** Herceptin PG 10% 4C H1

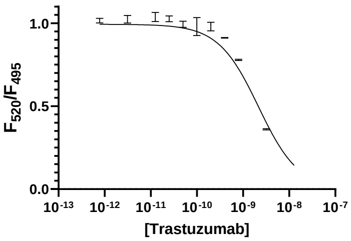


| Best-fit values | |
|---|---|
| Kd | = 1.248e-011 |
| Ki | 1.342e-011 |
| L | = 2.000e-009 |
| R | = 1.473e-010 |
| IC50 | 2.587e-009 |

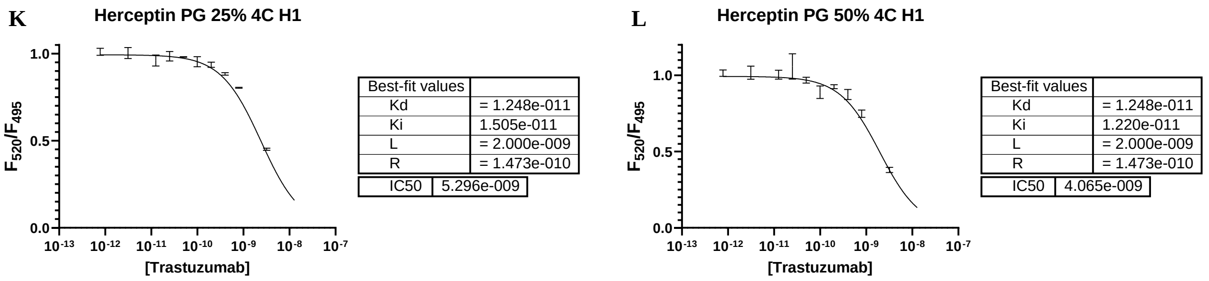


*Figure S29:* Effect of 2 h exposure to organic solvent on the apparent KI of trastuzumab in 1X PBS at room temperature. (A) Fab'-AF488 (B) Control (C) 10% DMF (D) 25% DMF (E) 50% DMF (F) 10% DMSO (G) 25% DMSO (H) 50% DMSO (J) 10% PG (K) 25% PG (L) 50% PG

#### B. Effect of Freeze/Thaw Cycles on the Binding Affinity of Trastuzumab

Aliquots of 1 mg trastuzumab in 200 µL 1X PBS at pH 7.4 were placed into microcentrifuge tubes and frozen at -20ºC or -80ºC for 2 h. After all aliquots were frozen, one sample from each freezer was labeled and placed into a storage compartment at -80ºC. The remaining samples were incubated at room temperature until thawed by inspection. This was repeated 10X, and samples were taken corresponding to 1X, 5X, and 10X freeze/thaw cycles. Following the protocol in Supplementary Section 2E, competition assays were equilibrated for 24 h at 4ºC with test wells containing 2 nM Fab'-AF488 and 500 pM HER2-ECD antigen. Curve-fitting was performed according to the general protocol in Supplementary Section 2F. The results are summarized in Figure S30. The dose-response curves and fits are shown in Figure S31.

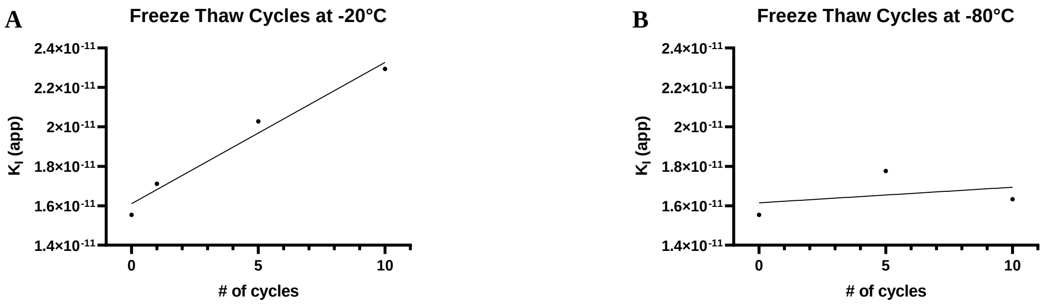


*Figure S30:* Effect of 0-10X freeze-thaw cycles on the apparent KI of trastuzumab in 1X PBS at pH 7.4 at (A) -20°C and (B) -80°C

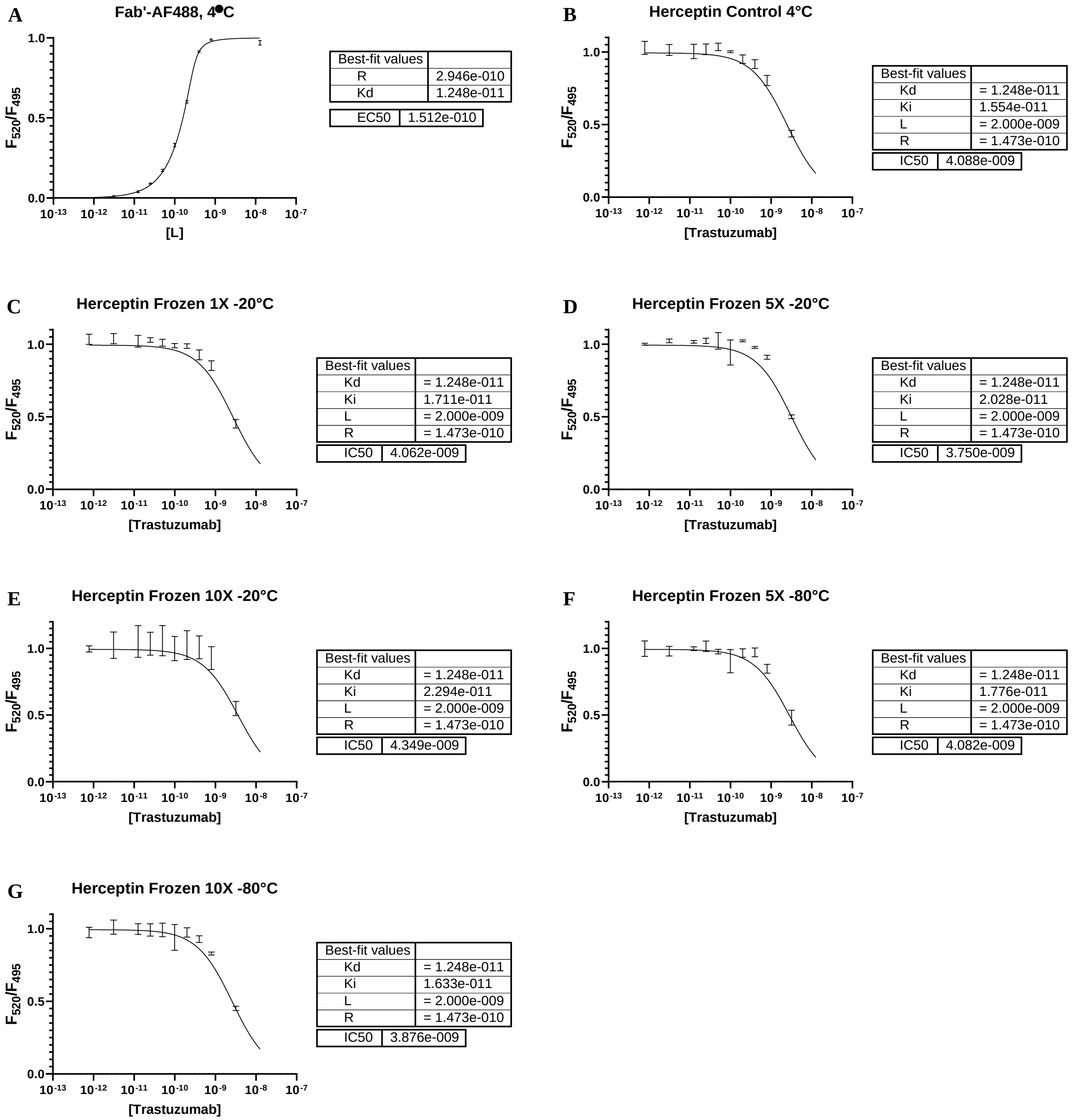


*Figure S31:* Effect of 0-10X freeze-thaw cycles on the apparent KI of trastuzumab in 1X PBS at pH 7.4. (A) Fab'-AF488 (B) Control (C,D,E) -20ºC (F,G) -80ºC

#### C. Effect of TCEP Reduction on the Binding Affinity of Trastuzumab

Trastuzumab (21 mg/mL) was exchanged into borate buffer (25 mM sodium borate, 25 mM NaCl, 1 mM DTPA, pH 8.0) with a Zeba™ spin column and adjusted to a concentration of 10 mg/mL. Aliquots of 2 mg were incubated with 0, 2, 4, 8, and 32 equivalents TCEP at 37ºC for 1.5 h with mild shaking. The samples were then buffer exchanged into 1X PBS on a Zeba™ spin column, and trastuzumab concentrations were determined by absorbance at 280 nm (ε=2.25e5 M-1cm-1.) The samples were frozen in liquid N2 and stored at -80ºC for further analysis. Competition assays were equilibrated for 18 h at 4ºC following the general protocol, with test wells containing 2 nM Fab'-AF488 and 500 pM HER2-ECD and data collected at intervals during the equilibration period. Curve-fitting was performed according to the general protocol in Supplementary Section 2F, and the $K_I$ values observed over the equilibration period were fit by a one-phase decay model to give the reported apparent $K_I$. The results are summarized in Figure S32. The one-phase decay model is shown in Figure S33. The dose-response curves and fits are shown in Figures S34-S39.

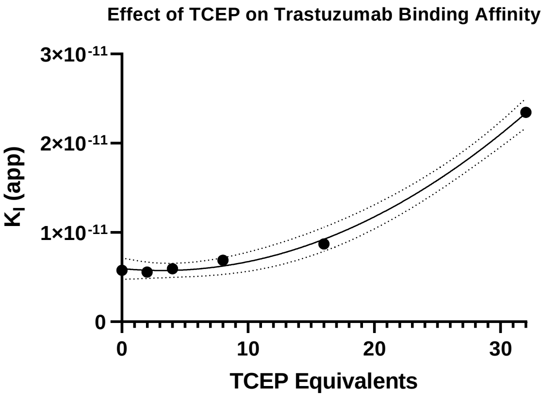


*Figure S33*: Effect of 1.5 h exposure to TCEP on the $K_I$ of trastuzumab in borate buffer (pH 8.0) at 37C.

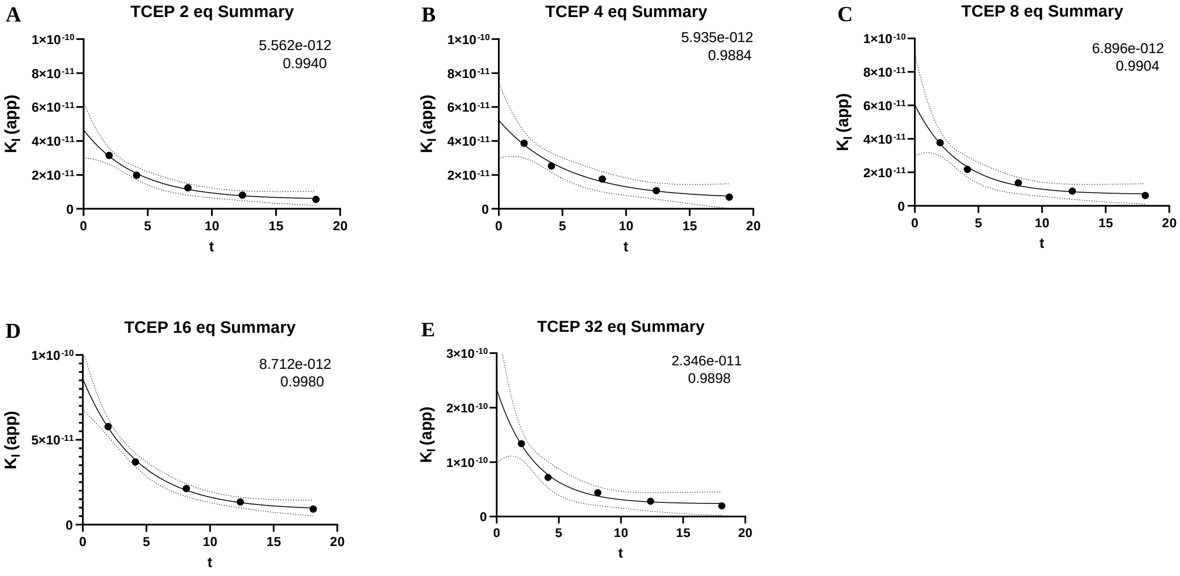


*Figure S34*: Effect of 1.5 h exposure to TCEP on the apparent KI of trastuzumab. Equilibration of apparent $K_I$ from 2-18 h. (A) 2 eq. (B) 4 eq. (C) 8 eq. (D) 16 eq. (E) 32 eq.

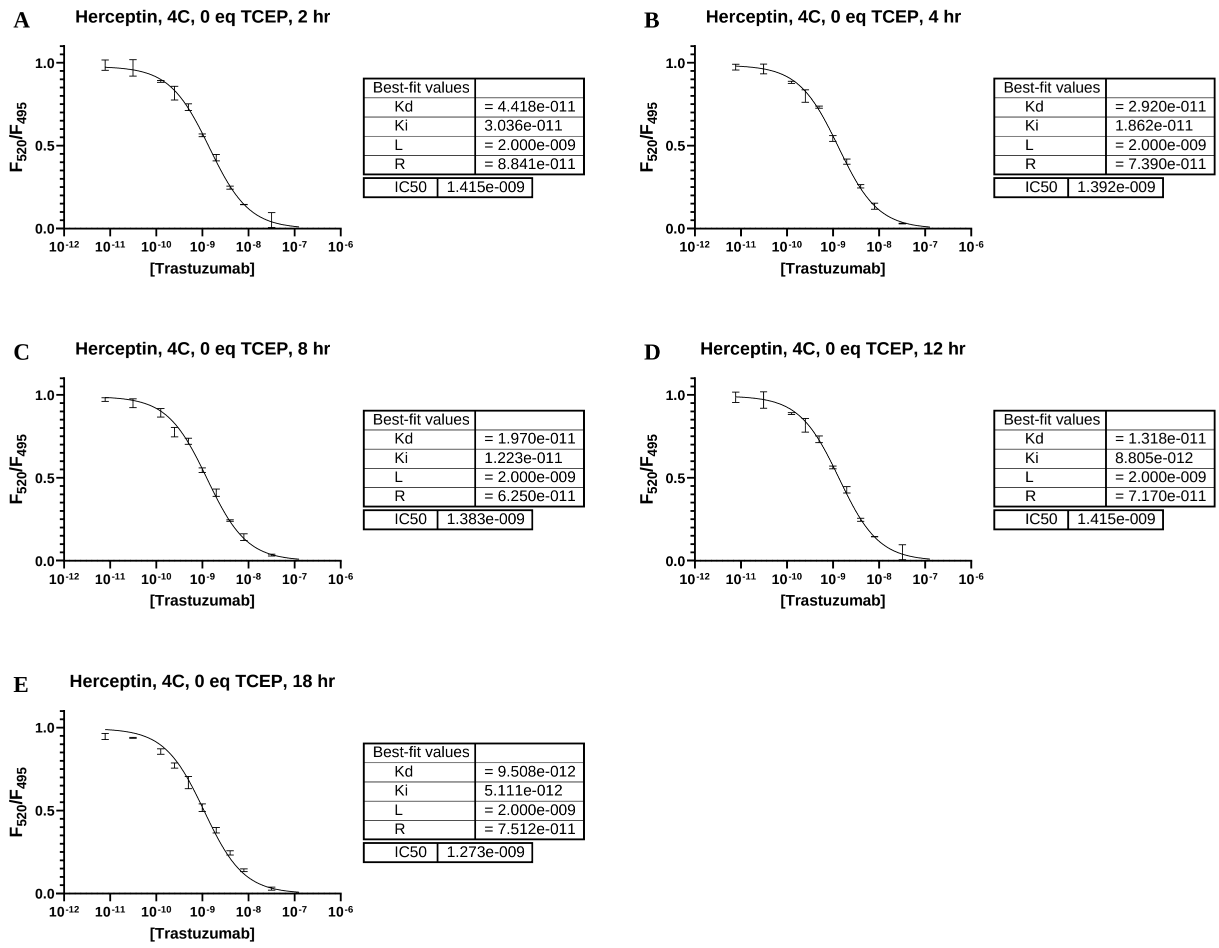


*Figure S35:* Effect of 1.5 h exposure to 0 eq. TCEP on the apparent $K_I$ of trastuzumab. Dose response curves and fitting of 2 nM Fab'-AF488 disassociation from HER2-ECD with increasing trastuzumab concentration at time points (A) 2h (B) 4h (C) 8h (D) 12h (E) 18h.

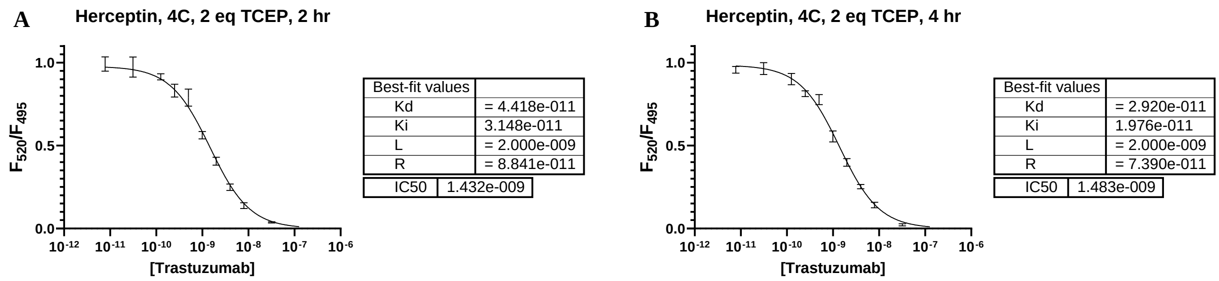

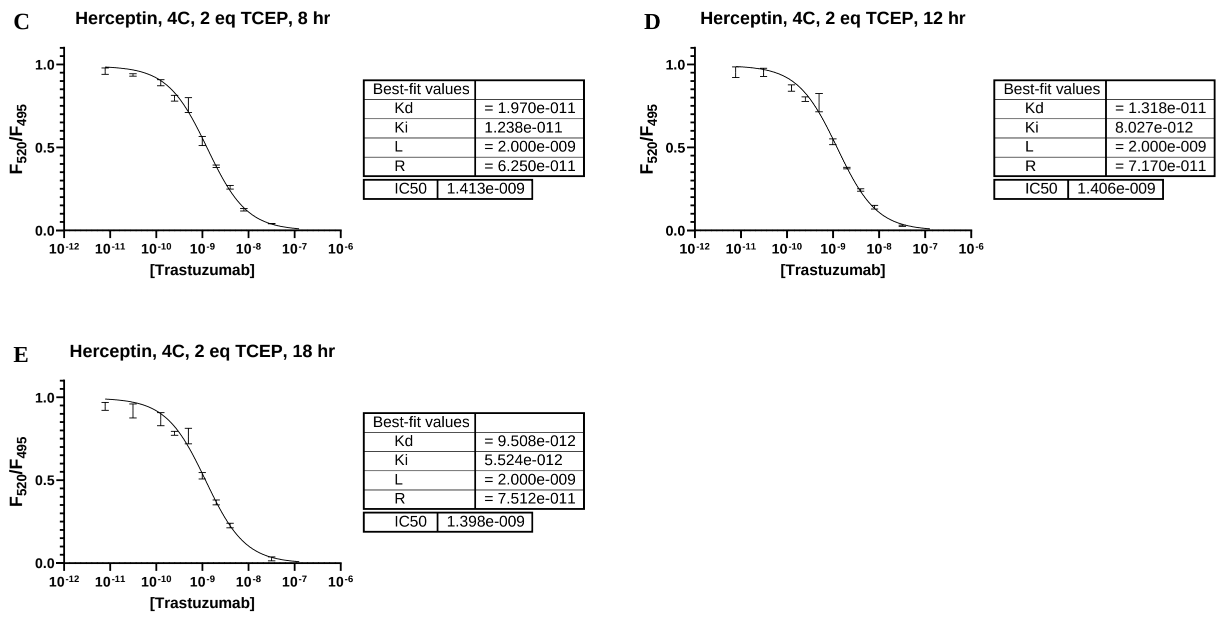


*Figure S36:* Effect of 1.5 h exposure to 2 eq. TCEP on the apparent $K_I$ of trastuzumab. Dose response curves and fitting of 2 nM Fab'-AF488 disassociation from HER2-ECD with increasing trastuzumab concentration at time points (A) 2h (B) 4h (C) 8h (D) 12h (E) 18h.

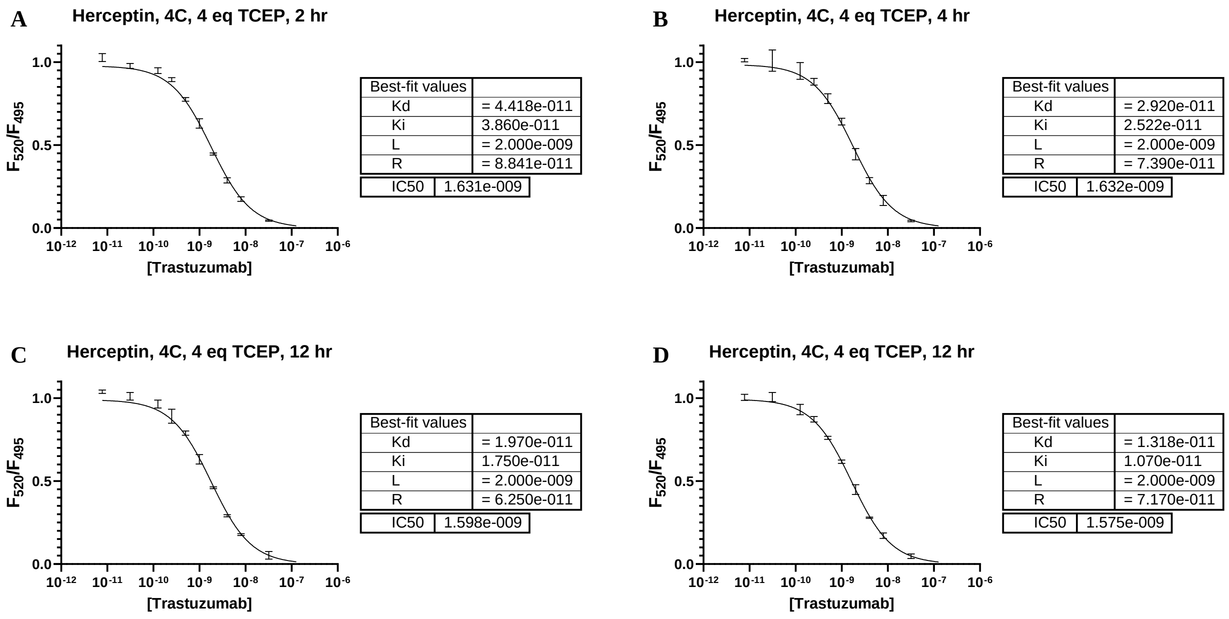

**E** Herceptin, 4C, 4 eq TCEP, 18 hr

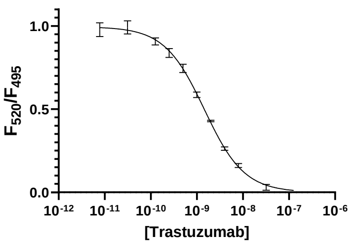


| Best-fit values | |
|---|---|
| Kd | = 9.508e-012 |
| Ki | 6.844e-012 |
| L | = 2.000e-009 |
| R | = 7.512e-011 |
| IC50 | 1.521e-009 |

*Figure S37:* Effect of 1.5 h exposure to 4 eq. TCEP on the apparent $K_I$ of trastuzumab. Dose response curves and fitting of 2 nM Fab'-AF488 disassociation from HER2-ECD with increasing trastuzumab concentration at time points (A) 2h (B) 4h (C) 8h (D) 12h (E) 18h.

**A** Herceptin, 4C, 8 eq TCEP, 2 hr

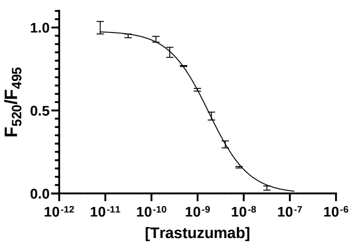


| Best-fit values | |
|---|---|
| Kd | = 4.418e-011 |
| Ki | 3.774e-011 |
| L | = 2.000e-009 |
| R | = 8.841e-011 |
| IC50 | 1.807e-009 |

**B** Herceptin, 4C, 8 eq TCEP, 4 hr

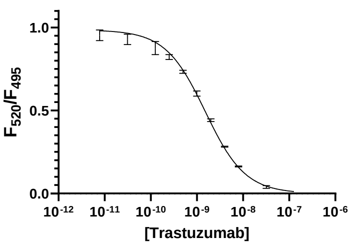


| Best-fit values | |
|---|---|
| Kd | = 2.920e-011 |
| Ki | 2.178e-011 |
| L | = 2.000e-009 |
| R | = 7.390e-011 |
| IC50 | 1.775e-009 |

**C** Herceptin, 4C, 8 eq TCEP, 8 hr

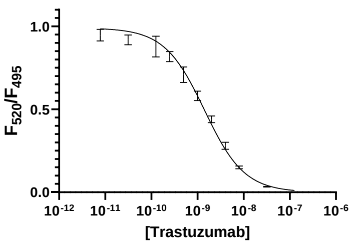


| Best-fit values | |
|---|---|
| Kd | = 1.970e-011 |
| Ki | 1.371e-011 |
| L | = 2.000e-009 |
| R | = 6.250e-011 |
| IC50 | 1.711e-009 |

**D** Herceptin, 4C, 8 eq TCEP, 12 hr

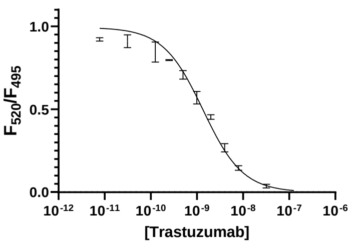


| Best-fit values | |
|---|---|
| Kd | = 1.318e-011 |
| Ki | 8.793e-012 |
| L | = 2.000e-009 |
| R | = 7.170e-011 |
| IC50 | 1.789e-009 |

**E** Herceptin, 4C, 8 eq TCEP, 18 hr

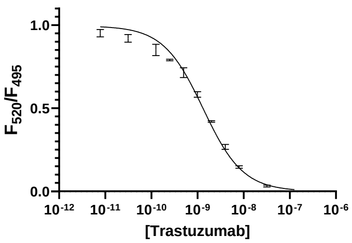


| Best-fit values | |
|---|---|
| Kd | = 9.508e-012 |
| Ki | 6.197e-012 |
| L | = 2.000e-009 |
| R | = 7.512e-011 |
| IC50 | 1.652e-009 |

*Figure S38:* Effect of 1.5 h exposure to 8 eq. TCEP on the apparent $K_I$ of trastuzumab. Dose response curves and fitting of 2 nM Fab'-AF488 disassociation from HER2-ECD with increasing trastuzumab concentration at time points (A) 2h (B) 4h (C) 8h (D) 12h (E) 18h.

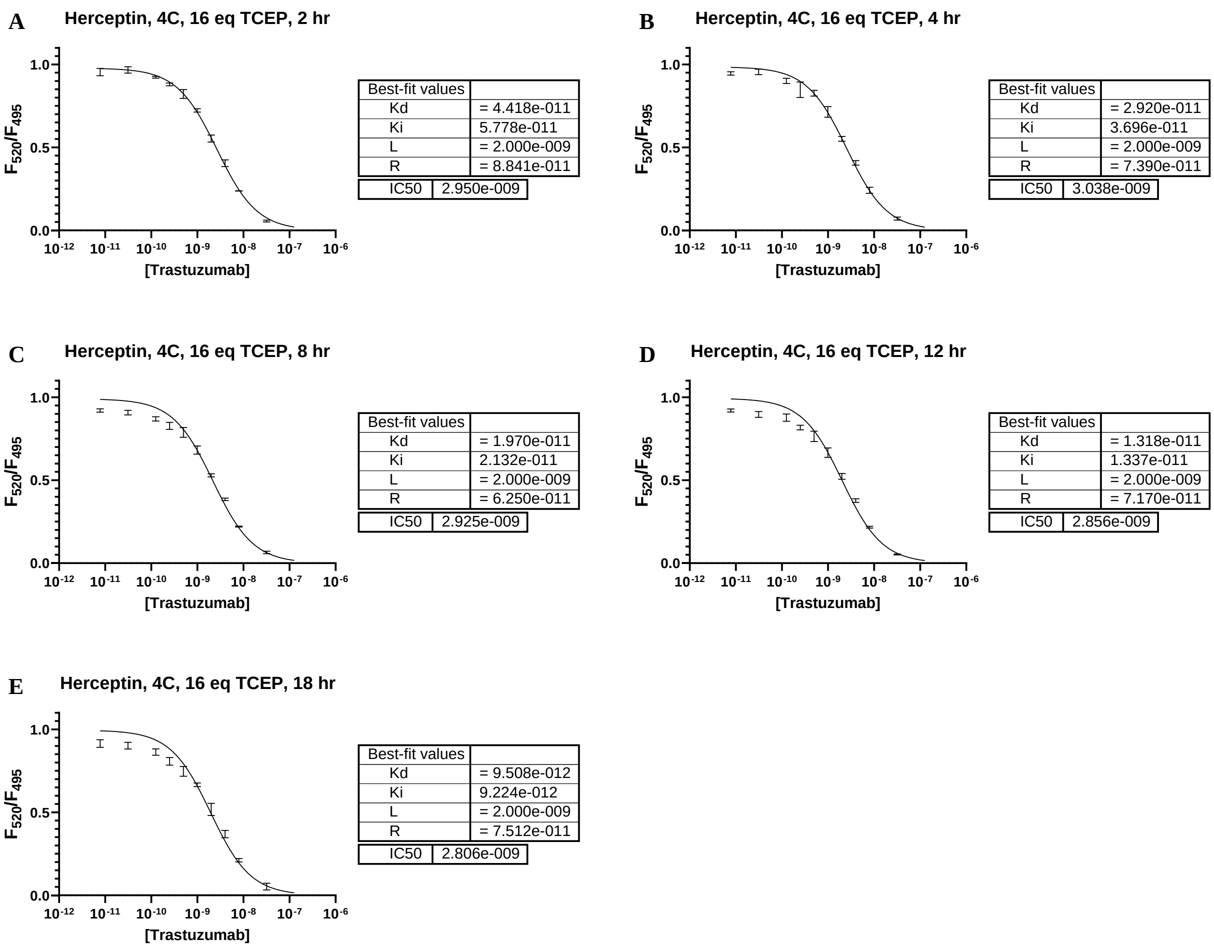


*Figure S39:* Effect of 1.5 h exposure to 16 eq. TCEP on the apparent $K_I$ of trastuzumab. Dose response curves and fitting of 2 nM Fab'-AF488 disassociation from HER2-ECD with increasing trastuzumab concentration at time points (A) 2h (B) 4h (C) 8h (D) 12h (E) 18h.

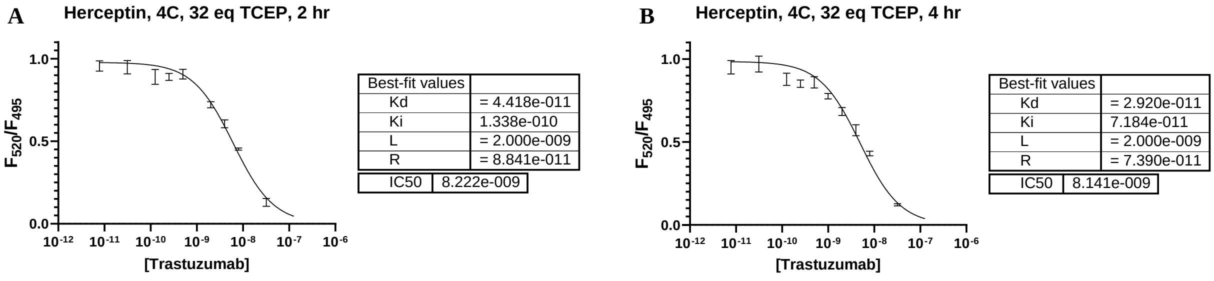

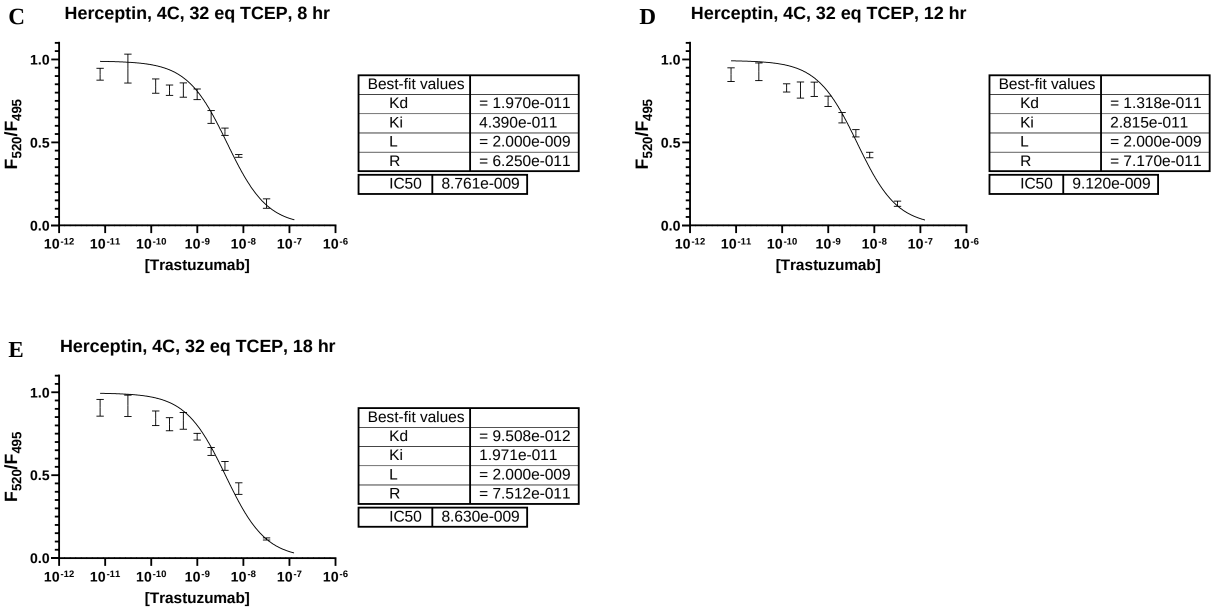


| Best-fit values (C) | |
|---|---|
| Kd | = 1.970e-011 |
| Ki | 4.390e-011 |
| L | = 2.000e-009 |
| R | = 6.250e-011 |
| IC50 | 8.761e-009 |

| Best-fit values (D) | |
|---|---|
| Kd | = 1.318e-011 |
| Ki | 2.815e-011 |
| L | = 2.000e-009 |
| R | = 7.170e-011 |
| IC50 | 9.120e-009 |

| Best-fit values (E) | |
|---|---|
| Kd | = 9.508e-012 |
| Ki | 1.971e-011 |
| L | = 2.000e-009 |
| R | = 7.512e-011 |
| IC50 | 8.630e-009 |

*Figure S40:* Effect of 1.5 h exposure to 32 eq. TCEP on the apparent $K_I$ of trastuzumab. Dose response curves and fitting of 2 nM Fab'-AF488 disassociation from HER2-ECD with increasing trastuzumab concentration at time points (A) 2h (B) 4h (C) 8h (D) 12h (E) 18h.

#### D. Synthesis and Effect of Stoichiometry on Apparent $K_I$ of Trastuzumab-DM1 ADCs

Trastuzumab (21 mg/mL) was exchanged into 1X PBS buffer with a Zeba™ spin column and adjusted to a concentration of 10 mg/mL. Aliquots of 2 mg trastuzumab in PBS were adjusted to 20% DMSO and incubated with 0, 8, 16, or 32 molar equivalents of DM-1 crosslinked with succinimidyl-4-[N-maleimidomethyl]cyclohexane-1-carboxylate (smcc-DM1) at 40ºC for 2 h with mild shaking. **Caution!** Cytotoxic drugs (smcc-DM1, 1228105-51-8) can be hazardous in small quantities and require standard personal protective equipment. The solution was cooled and the T-DM1 samples were separated from unconjugated drug-linker by dilution with 1X PBS on a 10K MWCO 0.5 mL Amicon® centrifugal filter, followed by gel filtration. The average DAR was calculated by the ratio of UV absorbance at 252 and 280 nm with equations outlined in Supplementary Section 3C. The average DAR of conjugates from 8, 16 and 32 equivalents was estimated by UV-Vis spectroscopy as 2.1, 3.2, and 2.7, respectively. To evaluate the effect of stoichiometry on the apparent $K_I$ of T-DM1, we performed a series of competition assays between these samples and 2 nM Fab'-AF488 with 500 pM HER2-ECD antigen. Following the protocol in Supplementary Section 2E/F, the assays were equilibrated for 18 h at 4ºC and the data collected at intervals during the equilibration. Curve-fitting was performed according to the general protocol in Supplementary Section 2F, and the $K_I$ values observed over the equilibration period were fit by a one-phase decay model to give the reported apparent $K_I$. The results are summarized in Figure S40. The one-phase decay models are shown in Figure S41. The dose-response curves and fits are shown in Figure S42-S44.

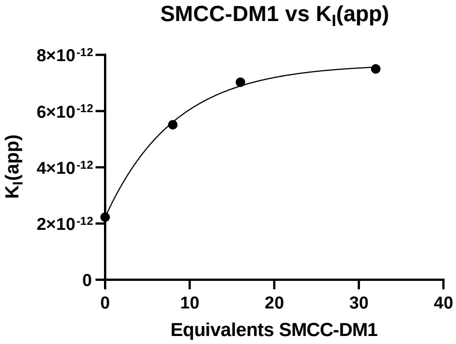


*Figure S40*: Effect of smcc-DM1 conjugation stoichiometry on the apparent KI of T-DM1 in 1X PBS.

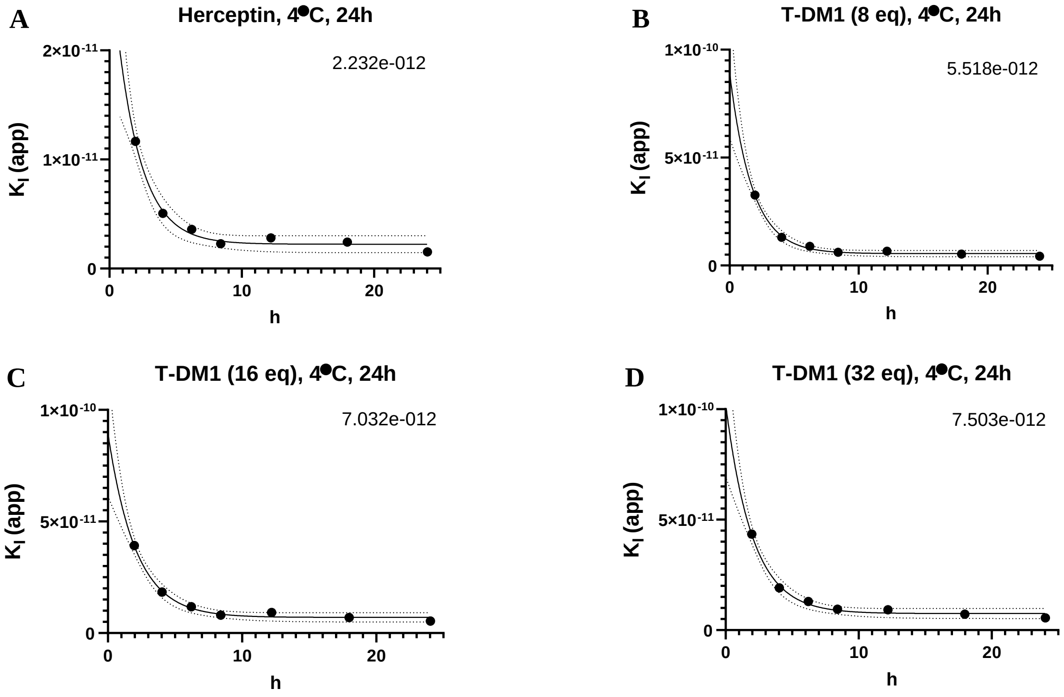


*Figure S41*: Effect of 2 h conjugation with smcc-DM1 at 40ºC on the apparent KI of trastuzumab. Equilibration of apparent KI from 2-24 h of conjugates produced by stoichiometry (A) 0 eq. (B) 8 eq. (C) 16 eq. (D) 32 eq.

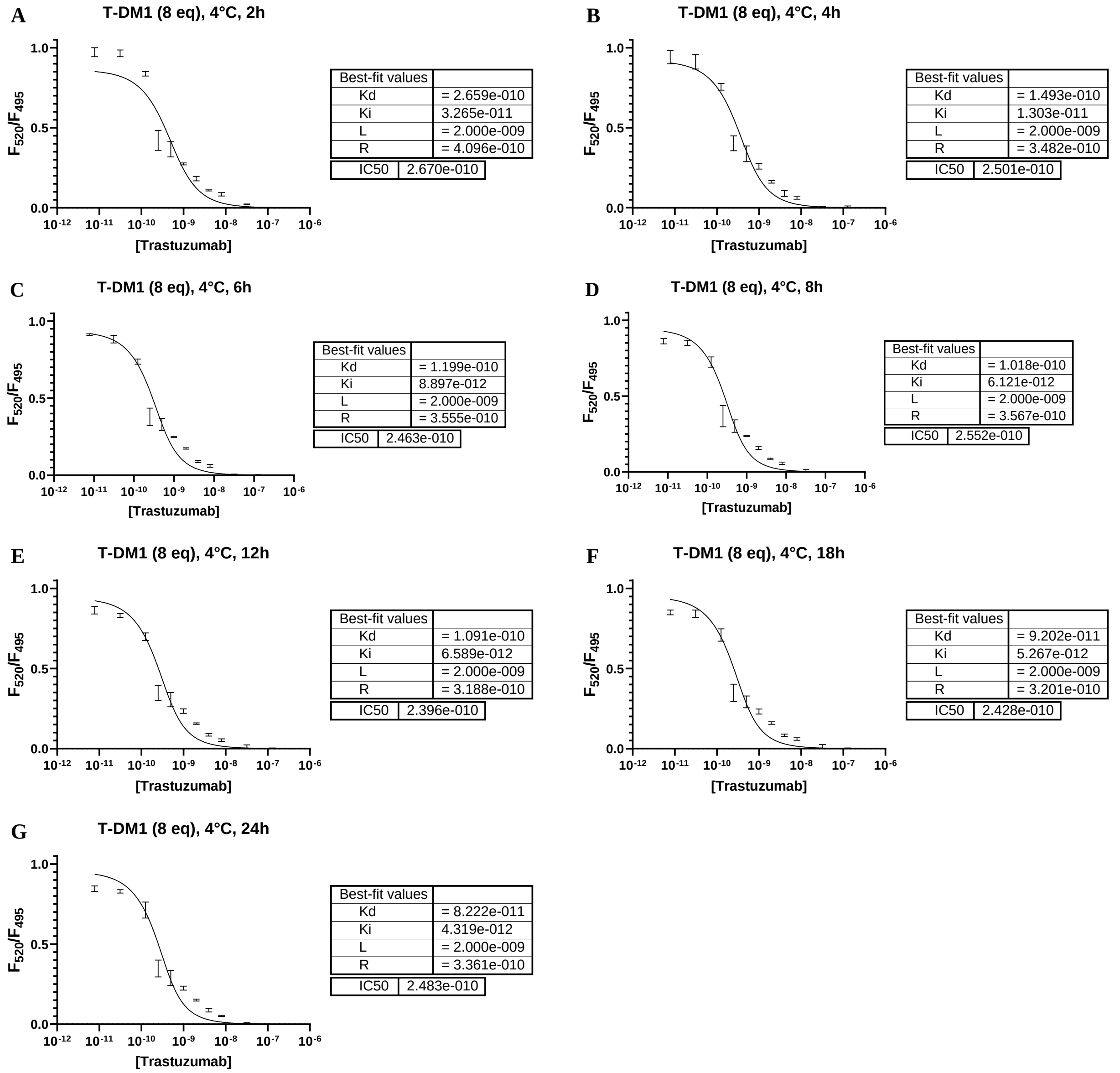


*Figure S42*: Effect of 2 h conjugation with smcc-DM1 at 40ºC on the apparent KI of trastuzumab. Equilibration of apparent $K_I$ from of T-DM1 conjugates produced with 8 eq. smcc-DM1 at (A) 2h (B) 4h (C) 6h (D) 8h (E) 12h (F) 18h (G) 24h

**A** T-DM1 (16 eq), 4°C, 2h

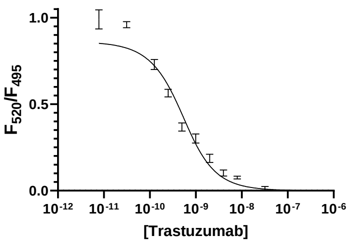


| Best-fit values | |
|---|---|
| Kd | = 2.659e-010 |
| Ki | 3.922e-011 |
| L | = 2.000e-009 |
| R | = 4.096e-010 |
| IC50 | 2.964e-010 |

**B** T-DM1 (16 eq), 4°C, 4h

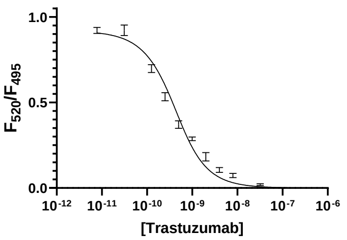


| Best-fit values | |
|---|---|
| Kd | = 1.493e-010 |
| Ki | 1.840e-011 |
| L | = 2.000e-009 |
| R | = 3.482e-010 |
| IC50 | 3.161e-010 |

**C** T-DM1 (16 eq), 4°C, 6h

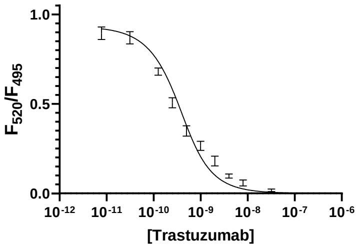


| Best-fit values | |
|---|---|
| Kd | = 1.199e-010 |
| Ki | 1.177e-011 |
| L | = 2.000e-009 |
| R | = 3.555e-010 |
| IC50 | 3.100e-010 |

**D** T-DM1 (16 eq), 4°C, 8h

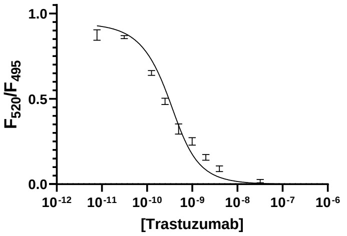


| Best-fit values | |
|---|---|
| Kd | = 1.018e-010 |
| Ki | 8.010e-012 |
| L | = 2.000e-009 |
| R | = 3.567e-010 |
| IC50 | 2.846e-010 |

**E** T-DM1 (16 eq), 4°C, 12h

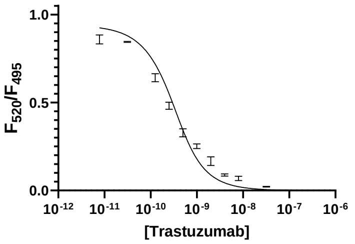


| Best-fit values | |
|---|---|
| Kd | = 1.091e-010 |
| Ki | 9.215e-012 |
| L | = 2.000e-009 |
| R | = 3.188e-010 |
| IC50 | 2.922e-010 |

**F** T-DM1 (16 eq), 4°C, 18h

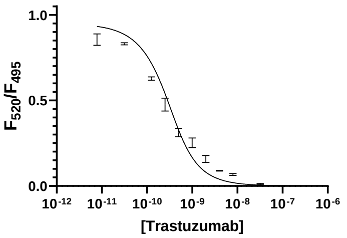


| Best-fit values | |
|---|---|
| Kd | = 9.202e-011 |
| Ki | 6.996e-012 |
| L | = 2.000e-009 |
| R | = 3.201e-010 |
| IC50 | 2.818e-010 |

**G** T-DM1 (16 eq), 4°C, 24h

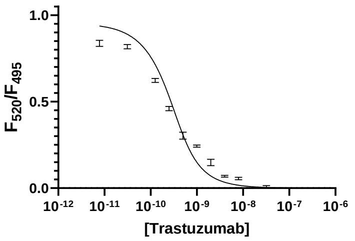


| Best-fit values | |
|---|---|
| Kd | = 8.222e-011 |
| Ki | 5.348e-012 |
| L | = 2.000e-009 |
| R | = 3.361e-010 |
| IC50 | 2.869e-010 |

*Figure S43*: Effect of 2 h conjugation with smcc-DM1 at 40ºC on the apparent KI of trastuzumab. Equilibration of apparent $K_I$ from of T-DM1 conjugates produced with 16 eq. smcc-DM1 at (A) 2h (B) 4h (C) 6h (D) 8h (E) 12h (F) 18h (G) 24h

**A** T-DM1 (32 eq), 4°C, 2h

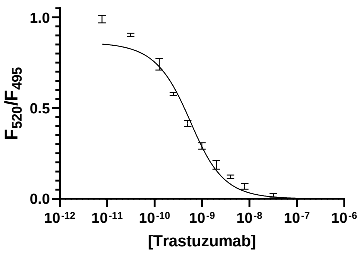


| Best-fit values | |
|---|---|
| Kd | = 2.659e-010 |
| Ki | 4.345e-011 |
| L | = 2.000e-009 |
| R | = 4.096e-010 |
| IC50 | 3.395e-010 |

**B** T-DM1 (32 eq), 4°C, 4h

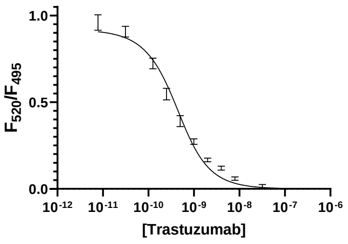


| Best-fit values | |
|---|---|
| Kd | = 1.493e-010 |
| Ki | 1.911e-011 |
| L | = 2.000e-009 |
| R | = 3.482e-010 |
| IC50 | 3.210e-010 |

C

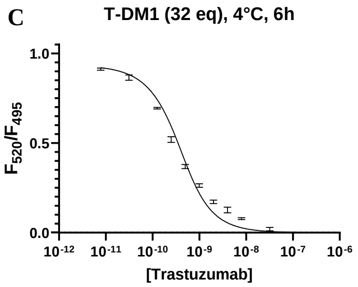


| Best-fit values | |
|---|---|
| Kd | = 1.199e-010 |
| Ki | 1.293e-011 |
| L | = 2.000e-009 |
| R | = 3.555e-010 |
| IC50 | 3.132e-010 |

D

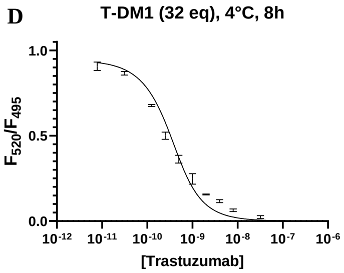


| Best-fit values | |
|---|---|
| Kd | = 1.018e-010 |
| Ki | 9.450e-012 |
| L | = 2.000e-009 |
| R | = 3.567e-010 |
| IC50 | 2.946e-010 |

E

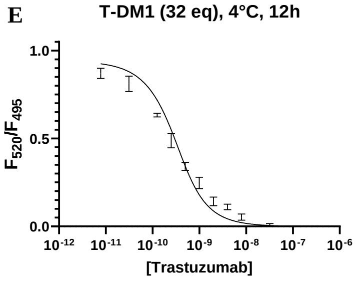


| Best-fit values | |
|---|---|
| Kd | = 1.091e-010 |
| Ki | 9.187e-012 |
| L | = 2.000e-009 |
| R | = 3.188e-010 |
| IC50 | 2.984e-010 |

F

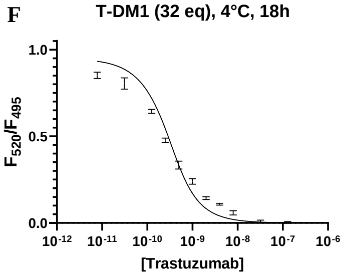


| Best-fit values | |
|---|---|
| Kd | = 9.202e-011 |
| Ki | 7.246e-012 |
| L | = 2.000e-009 |
| R | = 3.201e-010 |
| IC50 | 3.065e-010 |

G

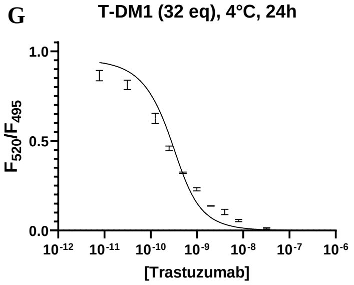


| Best-fit values | |
|---|---|
| Kd | = 8.222e-011 |
| Ki | 5.532e-012 |
| L | = 2.000e-009 |
| R | = 3.361e-010 |
| IC50 | 2.677e-010 |

*Figure S44*: Effect of 2 h conjugation with smcc-DM1 at 40ºC on the apparent KI of trastuzumab. Equilibration of apparent $K_I$ from of T-DM1 conjugates produced with 32 eq. smcc-DM1 at (A) 2h (B) 4h (C) 6h (D) 8h (E) 12h (F) 18h (G) 24h
